# A Novel Silver-Containing Antimicrobial potentiates aminoglycoside activity against *Pseudomonas aeruginosa*

**DOI:** 10.1101/2023.03.15.532855

**Authors:** Gracious Yoofi Donkor, Greg M. Anderson, Michael Stadler, Patrick Ofori Tawiah, Carl D. Orellano, Kevin A. Edwards, Jan-Ulrik Dahl

## Abstract

The rapid dissemination of antibiotic resistance combined with the decline in the discovery of novel antibiotics represents a major challenge for infectious disease control that can only be mitigated by investments into novel treatment strategies. Alternative antimicrobials, including silver, have regained interest due to their diverse mechanisms of inhibiting microbial growth. One such example is AGXX®, a broad-spectrum silver containing antimicrobial that produces highly cytotoxic reactive oxygen species (ROS) to inflict extensive macromolecular damage. Due to connections identified between ROS production and antibiotic lethality, we hypothesized that AGXX® could potentially increase the activity of conventional antibiotics. Using the gram-negative pathogen *Pseudomonas aeruginosa*, we screened possible synergistic effects of AGXX® on several antibiotic classes. We found that the combination of AGXX® and aminoglycosides tested at sublethal concentrations led to a rapid exponential decrease in bacterial survival and restored sensitivity of a kanamycin-resistant strain. ROS production contributes significantly to the bactericidal effects of AGXX®/aminoglycoside treatments, which is dependent on oxygen availability and can be reduced by the addition of ROS scavengers. Additionally, *P. aeruginosa* strains deficient in ROS detoxifying/repair genes were more susceptible to AGXX®/aminoglycoside treatment. We further demonstrate that this synergistic interaction was associated with significant increase in outer and inner membrane permeability, resulting in increased antibiotic influx. Our study also revealed that AGXX®/aminoglycoside-mediated killing requires an active proton motive force across the bacterial membrane. Overall, our findings provide an understanding of cellular targets that could be inhibited to increase the activity of conventional antimicrobials.

**IMPORTANCE:** The emergence of drug-resistant bacteria coupled with the decline in antibiotic development highlights the need for novel alternatives. Thus, new strategies aimed at repurposing conventional antibiotics have gained significant interest. The necessity of these interventions is evident especially in gram-negative pathogens as they are particularly difficult to treat due to their outer membrane. This study highlights the effectiveness of the silver containing antimicrobial AGXX® in potentiating aminoglycoside activities against *P. aeruginosa*. The combination of AGXX® and aminoglycosides not only reduces bacterial survival rapidly but also significantly re-sensitizes aminoglycoside-resistant *P. aeruginosa* strains. In combination with gentamicin, AGXX® induces increased endogenous oxidative stress, membrane damage and iron sulfur cluster disruption. These findings emphasize AGXX®’s potential as a route of antibiotic adjuvant development and shed light into potential targets to enhance aminoglycoside activity.

## INTRODUCTION

The spread of antibiotic resistance has now reached a large number of bacterial pathogens and evolved into a pertinent global health challenge (*1*). Over the past decade, resistance has been reported against all classical antibiotics, including last resort treatments such as polymyxins (*2*). The resistance crisis is further exacerbated in gram-negative pathogens due to the low permeability of their outer membrane and the extensive arsenal of drug resistance mechanisms that these critters employ. One example of a difficult to treat gram-negative bacterium is the opportunistic pathogen *P. aeruginosa*, a common cause of acute (e.g., wounds, burns) and chronic infections (e.g., diabetic ulcers, cystic fibrosis) (*3*). *P. aeruginosa* is characterized by its high intrinsic and acquired resistance mechanisms, allowing the pathogen to thrive in the presence of a large number of antibiotics (*4*).

In light of these challenges, recent studies have focused on alternative treatment strategies to limit bacterial infection and colonization (*5, 6*). Transition metals, such as silver and copper, have long been recognized for their antimicrobial activities and were already used by the ancient Greeks for would healing (*7*). Despite its long-standing history and high efficiency against bacteria, the antimicrobial mode of action of silver is poorly understood. Pleiotropic effects have been described and include changes in DNA condensation, membrane alteration and protein damage (*7, 8*). Silver ions have a particularly high affinity for cysteine thiols, disrupt exposed iron-sulfur clusters of dehydratase enzymes, and replace metal-containing cofactors, thus affecting a wide range of cytoplasmic and membrane proteins (*9, 10*). More recently, silver derivates have received increased attention in medical applications, e.g. as antimicrobial surface-coatings on catheters and implants to protect against biofilm-forming bacteria and to reduce the risk of nosocomial infections (*5, 11*). One such promising silver-based antimicrobial with broad-spectrum activity is AGXX®. AGXX® consists of micro-galvanic elements of silver (Ag) and ruthenium (Ru), which are surface conditioned with ascorbic acid (*12, 13*). AGXX® has been used in the form of a powder and implemented as an electroporated sheet on a variety of surfaces, including steel meshes, ceramics and water pipes, to limit bacterial colonization. AGXX®’s antimicrobial is not entirely dependent on the release of silver ions; instead it is proposed to generate reactive oxygen species (ROS), such as hydroxyl radicals (•OH) and superoxide (O_2_^-^) through a series of redox reactions where the oxidized Ag component is reduced by organic matter and donates electrons to the valent Ru, which subsequently generates O_2_^-^ and other ROS (*12–16*). Previous studies revealed that, compared to classical silver and other metals, AGXX® was significantly more bactericidal against gram-positive bacteria such as *Staphylococcus aureus* and *Enterococcus faecalis* (*12, 13*). Surprisingly, the antimicrobial effects of AGXX® on gram-negative pathogens remain largely unexplored.

Compounds that generate ROS and/or stimulate endogenous oxidative stress in bacteria have gained interest as potential antibiotic adjuvants (*17*). Adjuvants increase the efficacy of antibiotics by targeting metabolic processes or cellular networks that ultimately lead to a synergistic increase in antibiotic potency (*18*). A classic example for the synergy between adjuvants and antibiotic is the combination of amoxicillin, a beta-lactam antibiotic, and clavulanic acid. Due to clavulanic acid’s high affinity for beta-lactamase enzymes, its combination with the beta-lactam antibiotics, such as amoxicillin, synergistically potentiate their activity against penicillin-resistant bacteria (*19*). The hypothesis behind the synergistic effects of ROS-generating or -inducing compounds is based on the multimodal action of these compounds, which could potentially disrupt bacterial targets necessary for the defense of antibiotics (*20*). Recent studies on the bactericidal mode of action of aminoglycoside, fluoroquinolone and beta-lactam antibiotics have also proposed increased endogenous ROS stress as an additional mechanism for bacterial killing (*5, 20, 21*). Notably, activation of the bacterial envelope stress response, hyperactivation of the electron transport chain, and damage of iron sulfur clusters have been shown to contribute to an increase in endogenous ROS level (*5*). Although these findings are controversial, extensive evidence has been presented for the integral role of ROS-mediated damage in antibiotic-induced cell death (*6*). More importantly, recent studies employing ROS-generating compounds have reported promising results on their potential in sensitizing a wide variety of multidrug-resistant bacteria to both bactericidal and bacteriostatic antibiotics (*5*).

Given that AGXX®’s proposed mode of action involves ROS production and studies with focus on evaluating potential synergistic effects of AGXX® on conventional antibiotics are lacking, we started to investigate possible potentiating effects of AGXX® on members of several antibiotic classes, using the *P. aeruginosa* strain PA14 as a model. Exploring potential synergies between AGXX® and antibiotics could provide viable answers to the antimicrobial discovery drought for instance by reducing the minimal inhibitory concentrations of conventional antibiotics required for treatment or possibly overriding and/or delaying antibiotic resistance development (*22*). By exposing PA14 to sublethal concentrations of AGXX® and the antibiotics of interest alone and in combination, we found that bacterial survival was exponentially reduced when the cells were exposed to combinations of AGXX® and aminoglycoside antibiotics. Moreover, combined treatment of both compounds re-sensitized a kanamycin resistant PA14 strain to sublethal concentrations of the antibiotic. We further demonstrate that the combined treatment of AGXX® and aminoglycosides resulted in increased endogenous oxidative stress and a subsequent disruption of iron homeostasis, potentially providing an explanation for the elevated ROS level. The synergy was associated with a significant increase in outer and inner membrane permeability, which facilitates antibiotic influx. Moreover, our studies revealed that the synergy between AGXX® and aminoglycosides relies on an active proton motive force across the bacterial membrane.

## RESULTS

### AGXX® is more efficient in killing P. aeruginosa than silverdene and silver nitrate

Studies with silver ions suggest that gram-negative bacteria are more susceptible than gram-positive bacteria (*23*). In this study, we investigated the effect of AGXX® on *P. aeruginosa* using PA14 as model strain. To compare the effective antimicrobial concentrations of different AGXX® formulations against PA14, we first performed survival analyses in the presence and absence of AGXX383®, AGXX394®, AGXX823®, and AGXX720C®, respectively. While these AGXX® formulations all consist of the galvanized silver/ruthenium complex, they differ in various aspects such as silver ratio, particle size, and production procedure, which may affect their antimicrobial activity. Our time killing assays revealed that AGXX383® and AGXX394® have comparable antimicrobial activities against PA14 and are considerably more potent in inhibiting PA14 growth and survival compared to AGXX823® and AGXX720C® (**Supplementary FIG S1**). Silver derivates have received increased attention in medical applications, e.g. as antimicrobial surface-coatings on catheters that protect from biofilm-forming bacteria and reduce the risk of nosocomial infections (*24*). Moreover, silver is used in topicals to prevent and/or treat infections in wounds (*25*). One such example is silver sulfadiazine (silverdene), the gold standard for treating and preventing *P. aeruginosa* infections in burn wound patients (*26*). However, silverdene is associated with complications such as allergic reactions to the sulfadiazine moiety emphasizing the need for novel treatment therapies (*26*). To compare the antimicrobial activities of AGXX394®, silver nitrate (AgNO_3_) and silverdene, we conducted survival studies of PA14 in the presence and absence of 25 μg/ml of each compound. Survival of PA14 was highly compromised when cells were exposed to AGXX394® (FIG.1). A direct comparison of the killing efficiencies between AGXX394 and silverdene and silver nitrate after 3 hours of treatment revealed a ∼1-log and 2-log higher bactericidal activity for AGXX394®, respectively, potentially making AGXX® an attractive treatment/prevention alternative (FIG. 1). AGXX treatment has been shown to elicit oxidative stress responses shock responses and other general stress response regulons (*13*). To confirm ROS production of AGXX394®-treated PA14, we quantified intracellular ROS levels using the redox-sensitive molecular probe dichlorodihydrofluorescein diacetate (H_2_DCFDA). H_2_DCFDA has been extensively demonstrated to detect various ROS, including peroxides as well as peroxyl- and •OH radicals (*27*). When exposed to ROS, the nonfluorescent fluorescent H_2_DCFDA, is oxidized to highly green fluorescent 2’,7’-dichlorofluorescein (DCF) (*27*). We found that exposure to both sublethal (20 µg/ml) and bactericidal (40 µg/ml) levels of AGXX394® caused a 2.5 and 4-fold increase in endogenous ROS levels **(Supplementary FIG S2A)**. Pretreating PA14 with thiourea, an ROS scavenger, resulted in reduced ROS levels and recovered survival at bactericidal AGXX394® concentrations (40 µg/ml) **(Supplementary FIG S2A&B)**.

**FIG 1.**
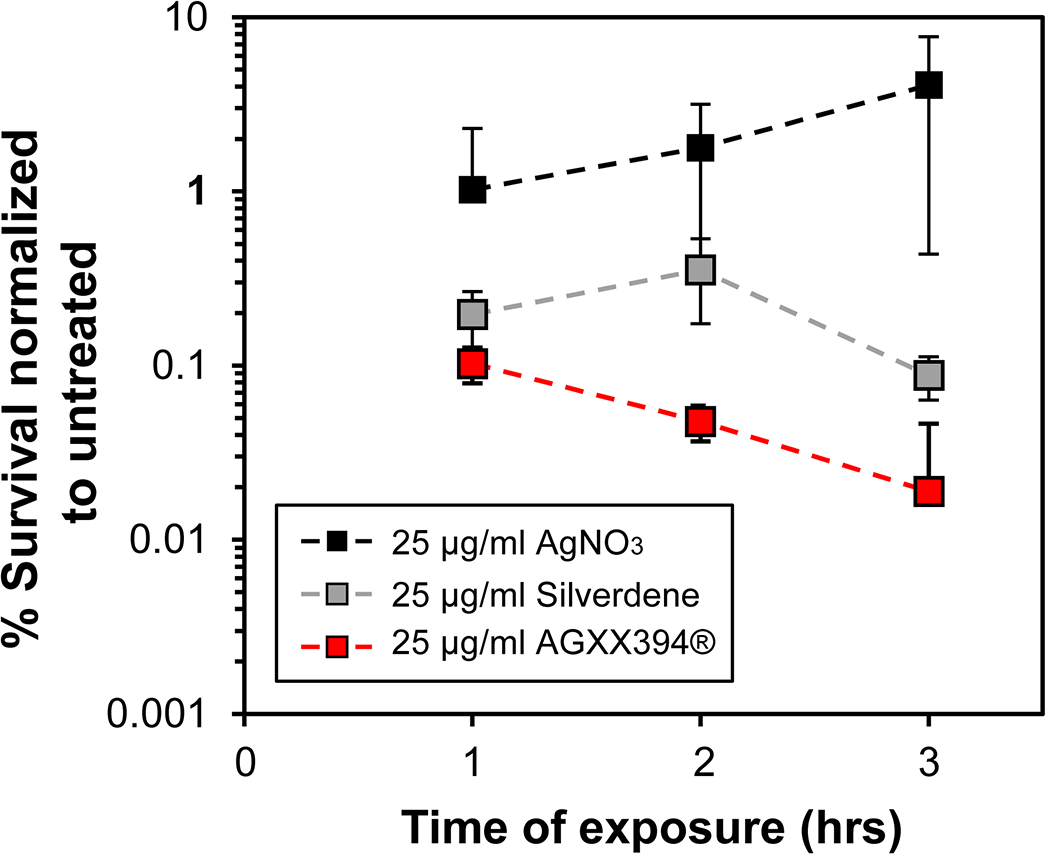
AGXX394® is more efficient in killing *P. aeruginosa* than silverdene and silver nitrate. Overnight PA14 cultures were diluted ∼25-fold into MOPSg media (OD_600_ = 0.1) and treated for three hours with 25 µg/ml AgNO_3_ (black square), silverdene (grey square), and AGXX394® (red square), respectively. Colony survival was evaluated every hour by serially diluting samples and plating them onto LB agar. Percent survival was calculated relative to the untreated control (*n=4, ± S.D)*.

### AGXX® exponentially increases the activity of aminoglycosides against P. aeruginosa

Our next goal was to determine how the presence of AGXX would affect antibiotic activity against *P. aeruginosa.* Specifically, we were interested in determining whether combining AGXX® with members of different antibiotic classes, such as inhibitors of DNA replication (fluoroquinolones), cell wall biosynthesis (β-lactams), folate biosynthesis, membrane integrity (polymyxins), and translation (aminoglycosides), show increased bactericidal activity against *P. aeruginosa*. Here, we used AGXX720C®, a formulation with significantly lower antimicrobial activity that allowed us to control dosage of AGXX® more reliably (**Supplementary FIG S1**). We determined the minimal inhibitory concentrations (MIC) of AGXX720C® and ten members of five different antibiotic classes in PA14, which were cultivated under aerobic conditions in Mueller-Hinton broth (**Supplementary Table S1**). Using time killing assays, we exposed PA14 to sublethal concentrations of AGXX720C® (75 µg/ml), the indicated antibiotics, and their combinations at the same sublethal concentrations that were used for the individual treatments, respectively. We monitored colony forming units counts every 60 min over a time course of three hours (FIG 2; **Supplementary FIG S3**) and calculated the percent survival for each sample at the 3-hour time point relative to the untreated control (FIG 2). None of the individual treatments with either 75 µg/ml AGXX720C® or any of the antibiotics resulted in substantial killing of PA14 (FIG 2; **Supplementary FIG S3**). The combination of 75 µg/ml AGXX720C® with 35 ng/ml ciprofloxacin and 100 ng/ml norfloxacin, respectively, resulted in PA14 survival comparable to the individual treatments suggesting that AGXX® does not potentiate the bactericidal activities of fluoroquinolones against PA14 (FIG 2A; **Supplementary FIG S3A, B**). Likewise, the combined treatments of AGXX720C® and 78 µg/ml carbenicillin, 0.156 µg/ml imipenem, or 125 µg/ml trimethoprim did not significantly change PA14 survival compared to their individual treatments (FIG 2B, C; **Supplementary FIG S3C**). On the other hand, we found that the bactericidal activity of the membrane targeting antibiotic polymyxin B was increased by the presence of AGXX720C® as evidenced by a 1-log reduction in PA14 survival as compared to AGXX720C® or polymyxin B alone, which each caused less than 5% killing (FIG 2D; **Supplementary FIG S3D**). However, we observed the most drastic decrease in PA14 survival when AGXX720C® was combined with a member of aminoglycoside antibiotics, even at concentrations far below the MIC. Co-treatment of AGXX720C® with 0.4 µg/ml gentamicin (Gm) (0.2x MIC) reduced PA14 survival by as much as 4 logs after three hours of treatment (FIG 2E; **Supplementary FIG S3E**), while combinations of AGXX720C® with 2 µg/ml amikacin (0.27x MIC), 1 µg/ml tobramycin (0.4x MIC), or 3 µg/ml streptomycin (0.15x MIC) caused up to 3 log reduction in survival (FIG 2E, **Supplementary FIG S3F-H**).

**FIG 2.**
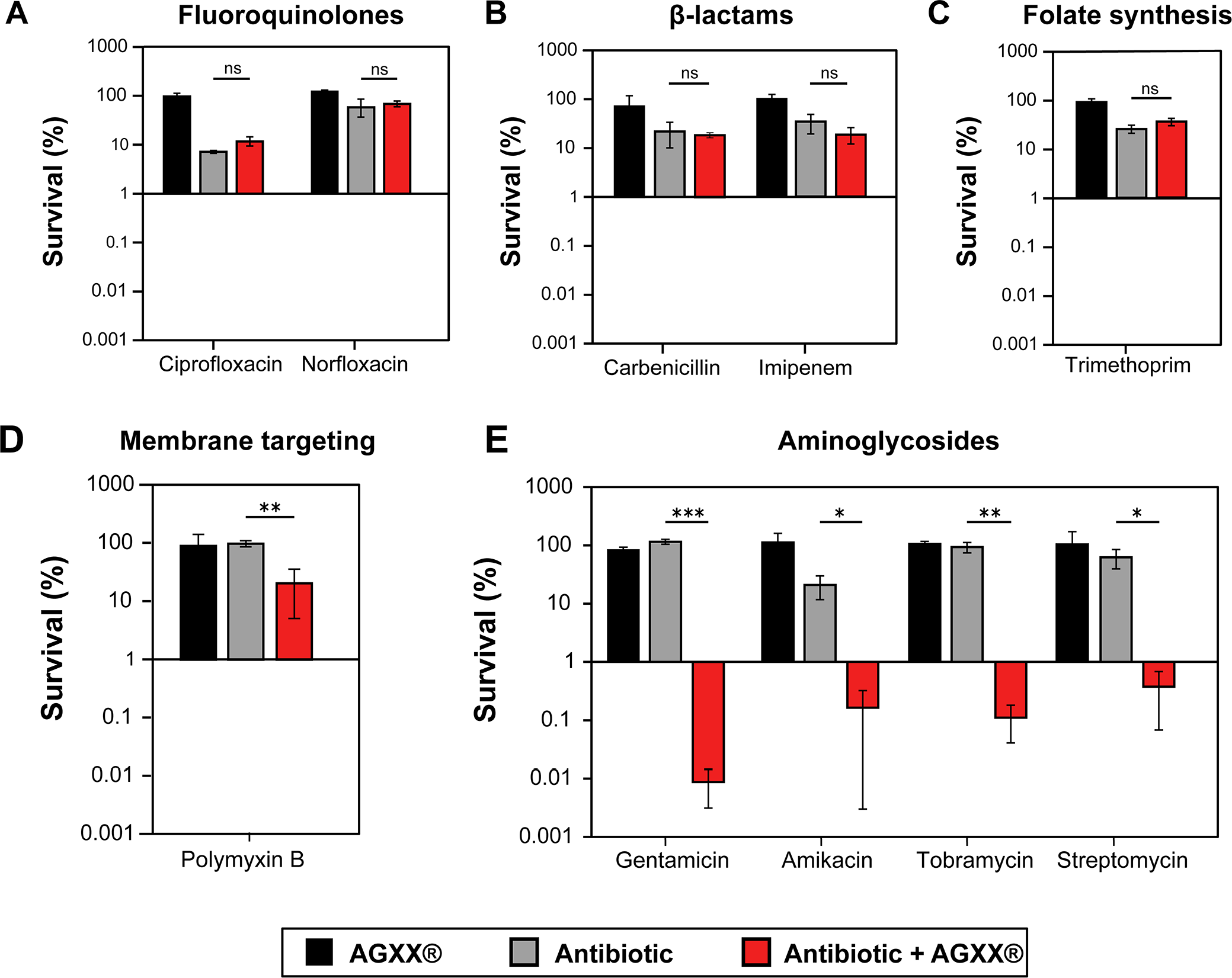
AGXX® exponentially increases the activity of aminoglycosides against *P. aeruginosa*. In our time killing assay, overnight PA14 cultures were diluted ∼25-fold (OD600 = 0.1) into MHB and exposed to 75 µg/ml AGXX720C® (black bars), a sublethal concentration of the indicated antibiotic (grey bar), or the combined treatment of both (red bars) for 3 hours. Samples were taking every 60 min, serial diluted, plated on LB agar, and incubated for 20 hours for CFU counts. Percent survival was calculated relative to the untreated control for: (A) 35 ng/ml ciprofloxacin and 100 ng/ml nalidixic acid, respectively; (B) 78 µg/ml carbenicillin and 0.156 µg/ml imipenem, respectively; (C) 125 µg/ml trimethoprim; (D) 1.5 µg/ml polymyxin B; and (E) 0.4 µg/ml gentamicin, 2.0 µg/ml amikacin, 1.0 µg/ml tobramycin, and 3 µg/ml streptomycin, respectively. All experiments were performed in at least three biological replicates and error bars represent mean (± SD. * p < 0.05, ** p < 0.01, *** p < 0.001; student’s t test, calculated relative to cultures treated with antibiotics alone).

It has been reported that AgNO_3_’s synergizing effects on aminoglycosides was reduced in minimal media (*28*). We therefore exposed PA14 sub-cultured in MOPS-glucose (MOPSg) minimal medium to increasing sublethal AGXX concentrations alone or in combination with a sublethal gentamicin **(Supplementary FIG S4A)**. Increasing AGXX® concentrations significantly reduced PA14 survival in a dose-response like manner **(Supplementary FIG S4A)**. Considering that casamino acid (CAS) supplementation was also reported to partially restore AgNO_3_’s effects in minimal media, we replicated our time killing assays in MOPSg media in the presence of 0.2% CAS. Surprisingly, addition of 0.2% CAS decreased AGXX®’s effects on Gm in MOPSg medium **(Supplementary FIG S4B)**. Although, we do not probe the reasons behind the effects of casamino acids in minimal media, it is evident that AGXX® synergizes with aminoglycosides in both rich and minimal media.

### AGXX® increases the sensitivity of P. aeruginosa strain PA14 to kanamycin

Considering the significant increase in lethality upon concurrent exposure of PA14 to AGXX720C® and aminoglycosides, we sought to determine whether the addition of AGXX720C® could reintroduce sensitivity in aminoglycoside-resistant *P. aeruginosa* strains. Like many other *P. aeruginosa* strains, PA14 is intrinsically resistant to many different antibiotics, including the aminoglycoside kanamycin. Under the conditions tested, our PA14 strain showed a MIC of 240 µg/ml for kanamycin. We then exposed PA14 to 75 µg/ml AGXX720C®, 50 µg/ml kanamycin, or a combination of 75 µg/ml AGXX720C® and 50 µg/ml kanamycin and determined their survival over the time course of 3 hours as described before. As early as 60 minutes post treatment, the combination of AGXX720C® and kanamycin reduced colony survival by ∼1.5 log, which increased to about 4 log (10,000-fold) difference in survival after 3 hours (FIG 3). In summary, we conclude from our data that AGXX® potentiates the activity of a wide range of aminoglycoside antibiotics, potentially making aminoglycoside-resistant *P. aeruginosa* strains more sensitive again.

**FIG 3.**
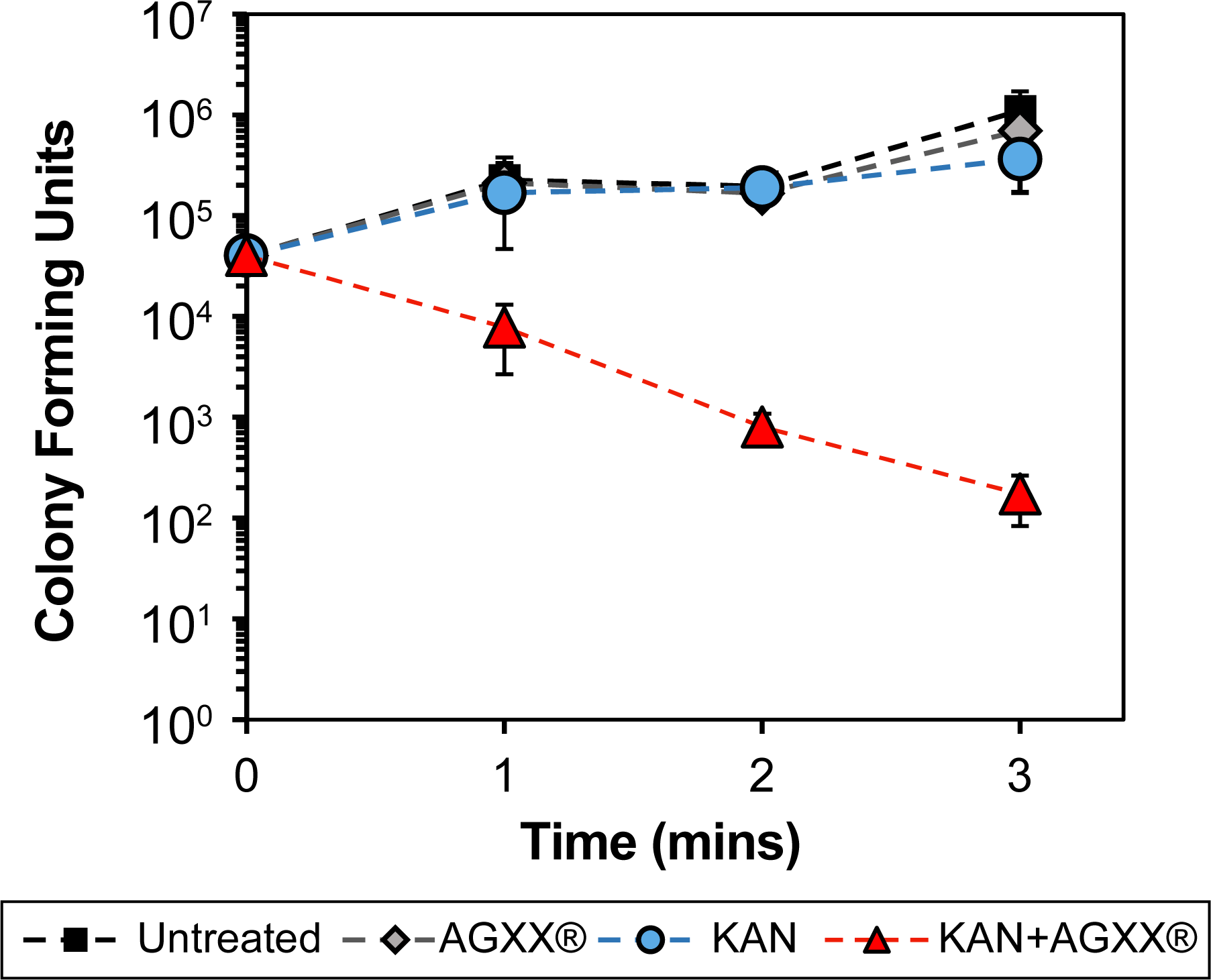
AGXX® increases the sensitivity of *P. aeruginosa* strain PA14 to kanamycin. Overnight PA14 cultures were diluted ∼25-fold into Mueller-Hinton media (OD_600_=0.1) and either left untreated or exposed to 75 μg/ml AGXX720C®, 50 μg/ml kanamycin, or the combination thereof for 3 hours. Samples were taken every 60 minutes, serial diluted and plated on LB agar for CFU counts, (*n=4, ± S.D)*.

### The combination of sublethal concentrations of AGXX® and aminoglycosides increases ROS formation and causes upregulation of genes indicative of DNA damage and protein aggregation

AGXX®’s main antimicrobial mode of action is mediated by the generation of ROS (*39*). Increased endogenous oxidative stress has also been linked to the lethality of aminoglycosides and other bactericidal antibiotics (*29, 30*). It may therefore be possible that a disruption in the redox balance during antibiotic exposure increases lethality by amplifying the adventitious generation of ROS by antibiotics. We treated exponentially growing PA14 for 60 min with sublethal concentrations of Gm (0.25 µg/ml), AGXX720C® (50 µg/ml), or their combination and compared H2DCFDA fluorescence in each sample to the untreated control. Individual treatments with either Gm or AGXX720C® did not result in a significant increase in H_2_DCFDA fluorescence, excluding the possibility that substantial amounts of ROS were formed at these sublethal concentrations (FIG 4A). However, when applied in combination, we observed an 8-fold increase in H_2_DCFDA fluorescence indicative of increased ROS production under these conditions. Pre-treatment of PA14 with the ROS scavenger thiourea (*31*) or the hydrogen peroxide (H_2_O_2_) -detoxifying enzyme catalase resulted in a significant decline in H_2_DCFDA fluorescence (FIG 4A) as well as restored PA14 survival by 2- to 4-log, respectively (FIG 4B; **Supplementary FIG S5A**). Thus, our data suggest that the high antimicrobial activity of the combinational treatment can at least in parts be explained by increased ROS formation. However, H_2_DCFDA is rather unspecific and detects numerous ROS compounds. We therefore followed up on our observation with the use of additional, more specific ROS-detecting fluorescent dyes. We used the boronate-based peroxy orange 1 (PO1) dye as well as hydroxyphenyl fluorescein (HPF) to examine intracellular concentrations of H_2_O_2_ and •OH levels, respectively (*32, 33*). After 1 hour of treatment, we detected a significant shift in PO1 and HPF fluorescence, indicating that both H_2_O_2_ and •OH are produced in AGXX720C®/Gm-treated PA14 (FIG 4C, D; **Supplementary FIG S5B, C**).

**FIG 4.**
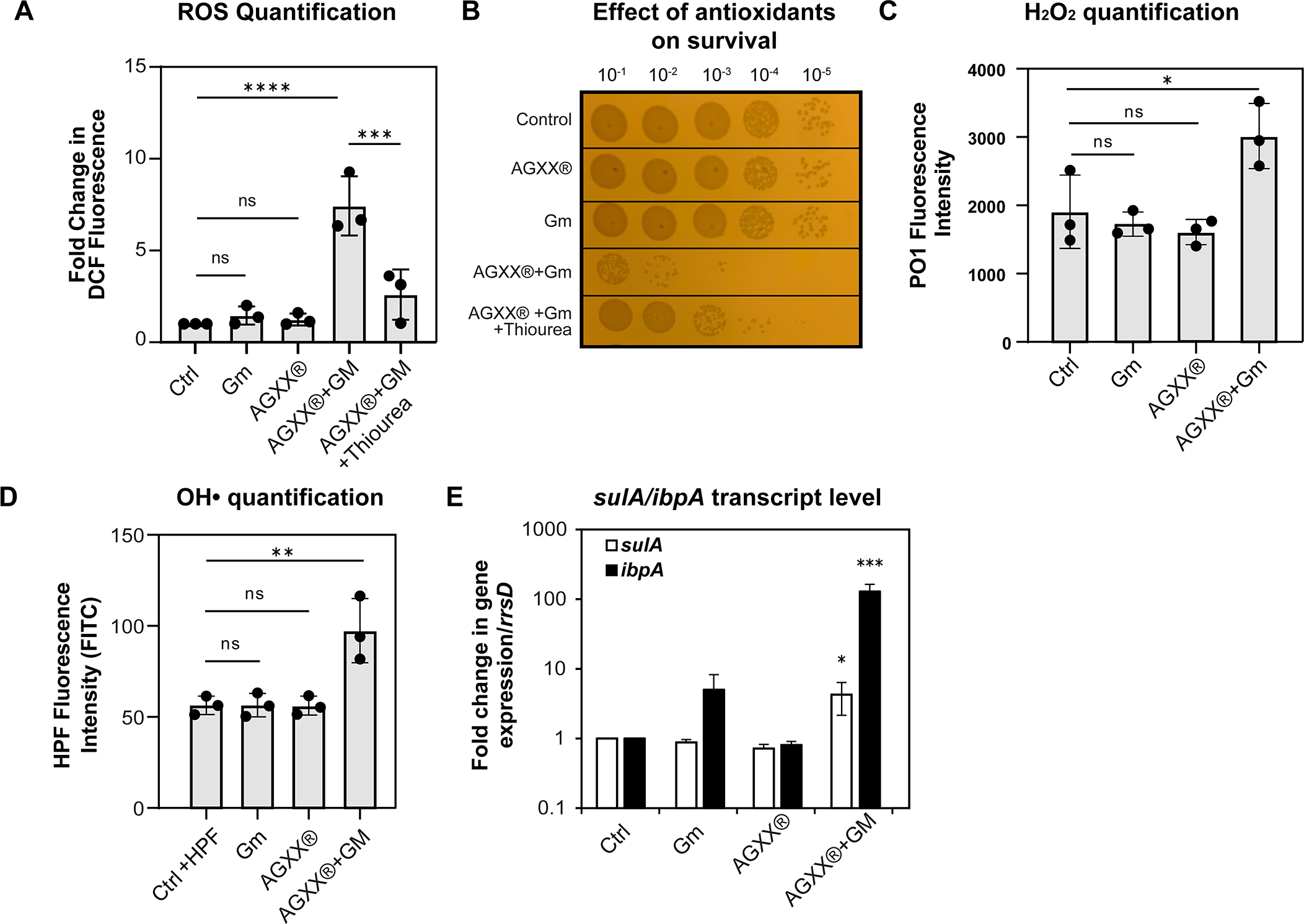
The combination of sublethal concentrations of AGXX® and gentamicin increases ROS levels and causes DNA damage and protein aggregation. Mid-log PA14 cells were treated with sublethal concentrations of Gm (0.25 µg/ml), AGXX720C® (50 µg/ml), the combination thereof, or left untreated. **(A)** Intracellular ROS levels were quantified by H_2_DCFDA fluorescence. 50mM thiourea was used as a ROS quencher (*n=3, ± S.D.)*. **(B)** Samples were serially diluted in PBS after 60 min of incubation, spot-titered onto LB agar and incubated for 20 hours. One representative of three independent experiments with similar outcomes. **(C, D)** Cells were strained with **(C)** 10 µM peroxy-orange 1 (PO1) and **(D)** 10µM hydroxyphenyl fluorescein (HPF) for 60 min and fluorescence was measured via flow cytometry (*n=3, ± S.D.)*. **(E)** The induction of *sulA* (white bar) and *ibpA* (black bar) transcript levels was determined by qRT-PCR. Gene expression was normalized to the housekeeping gene *rrsD* and calculated as fold changes based on expression levels in the untreated control (*n=3, ± S.D.*; One-way ANOVA, Dunnett’s posttest; *ns*=*p>0.05,* * *p* < 0.05, ** *p* < 0.01, *** *p* < 0.001, **** *p* < 0.0001).

To deal with the negative consequences of ROS and eliminate ROS-mediated damage, microorganisms have evolved intricate systems to maintain and restore a balanced redox homeostasis and repair ROS-mediated damage [recently reviewed in (*34–39*)]. Proteins and nucleic acids represent the most prevalent targets of ROS (*40, 41*). To test whether the elevated ROS production causes macromolecular damage, we analyzed the transcript level of *ibpA* and *sulA,* two genes that have previously been shown to be upregulated when cells experience severe oxidative stress (*42, 43*), in PA14 cells treated with Gm and AGXX alone and in combination. *ibpA* encodes a molecular chaperone, which plays a significant role in protecting bacteria from proteotoxic stressors including ROS (*44*). *sulA* encodes the cell division inhibitor SulA, which is part of the SOS response and induced when the cell experiences DNA damage, a possible consequence of oxidative stress (*45*). We exposed exponentially growing PA14 cells to sublethal concentrations of AGXX720C®, Gm, or the combination thereof for 60 min and quantified *ibpA/sulA* mRNA level. While individual treatments with sublethal concentrations of AGXX720C® or Gm did not cause significant changes in *ibpA/sulA* expression, their transcript levels were approximately 5-fold (*sulA*) and 100-fold (*ibpA*) increased when PA14 was treated with the combination of AGXX720C® and gentamicin, indicating significant macromolecular damage (FIG 4E).

### Anaerobic growth and antioxidant systems provide protection against ROS-mediated damage caused by a combinational treatment of AGXX® and aminoglycosides

To provide additional evidence that ROS contribute to the synergy between aminoglycosides and AGXX®, we determined whether their killing efficiency depends on the presence of molecular oxygen, which is essential for ROS production. PA14 was grown in MHB under either aerobic or anaerobic conditions and treated with sublethal concentrations of gentamicin, AGXX720C® or the combination thereof. When grown anaerobically, MHB was supplemented with 1% KNO_3_ to stimulate the proton motive force (PMF). Anaerobically respiring PA14 cells tolerated treatments with AGXX® and 0.5 µg/ml Gm (0.25x MIC) fairly well and showed about 2.5 log higher survival relative to aerobically grown cells exposed to the same treatment. However, when we increased the Gm concentration to 1 µg/ml (0.5x MIC) in the in the presence of AGXX®, PA14 survival also declined albeit at a lower rate than aerobically growing cells under the same conditions (FIG. 5A).

**FIG 5.**
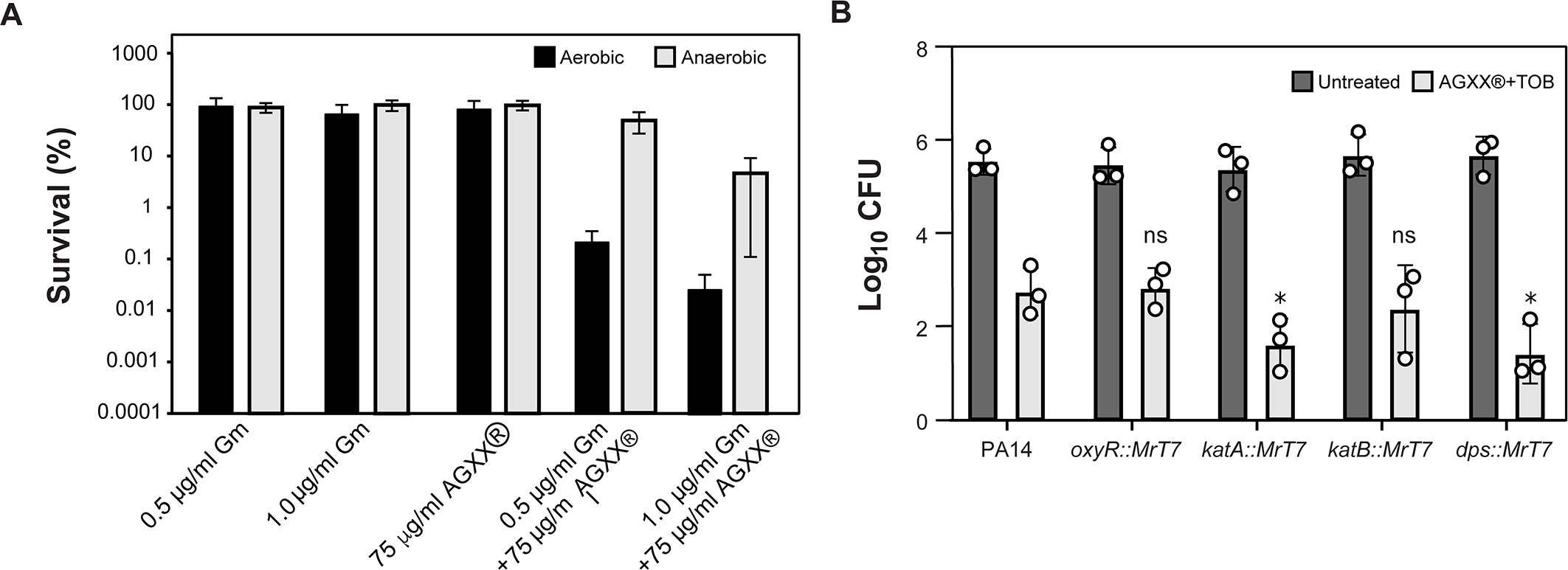
Anaerobic growth and antioxidant systems provide protection against ROS-mediated damage caused by a combinational treatment of AGXX® and aminoglycosides. **(A)** PA14 were diluted approximately 1000-fold (OD_600_=0.002) in MHB supplemented with 1% KNO_3_ and grown under aerobic (black bars) and anaerobic (grey bars) conditions, respectively. At OD_600_ ∼0.2, cultures were treated with either 0.5 µg/ml or 1.0 µg/ml Gm alone or in combination with 75 µg/ml AGXX® for four hours. Cells were subsequently plated onto LB agar for 20 hours at 37°C to enumerate surviving colonies (*n=3, ± S.D.*). **(B)** Overnight cultures of PA14 wildtype and mutant strains with MrT7 transposon insertions in *oxyR, katA, katB* and *dps* were diluted into MOPSg to an OD_600_=0.01 and grown under aerobic conditions until OD_600_=0.1. Cultures were either left untreated (grey bars) or treated with a combination of 0.25 µg/ml tobramycin and 25 µg/ml AGXX720C® (white bars). Bacterial survival was quantified after two hours by serially diluting cells in PBS and plating on LB agar for 20 hours at 37°C (*n=3, ± S.D*; Student t-test, * *p* < 0.05).

Next, we tested the susceptibility of PA14 strains with transposon insertions in genes that have previously been identified as major oxidative stress response and/or defense systems. We exposed PA14 transposon mutant strains (*46*) with defects in *oxyR* (encodes ROS-sensing transcriptional regulator OxyR), *katA* (encodes catalase KatA), *katB* (encodes catalase KatB), and *dps* (encodes an iron acquisition protein in response to oxidative stress) to sublethal concentrations of AGXX720C® (25 µg/ml), tobramycin (0.25 µg/ml), or the combination thereof for 3 hours and compared their colony survival to that of the corresponding wildtype strain PA14 (FIG. 5B). We found that strains deficient in functional copies of *katA* and *dps* showed a little over 1 log reduced survival compared to wildtype cells, while no difference in survival was observed for cells lacking the hydrogen peroxide global regulator OxyR and catalase KatB, respectively (FIG. 5B). Taken together, our findings point towards a relevant role for ROS generation in the synergistic interaction between aminoglycosides and AGXX®.

### The synergy between AGXX® and aminoglycosides on PA14 killing is in parts mediated by a disruption in iron homeostasis

Fe/S cluster are essential cofactors in various metabolic enzymes, including members of the electron transport chain and the tricarboxylic acid cycle (TCA) cycle (*47*). Fe/S cluster are particularly vulnerable to ROS, which negatively impacts the activity of metabolic enzymes and ultimately cellular metabolism during oxidative stress. Likewise, silver has been found to disrupt Fe/S clusters in proteins (*10*), resulting in a cellular increase in free iron, which in turn can stimulate •OH formation via Fenton reaction and cause extensive macromolecular damage (*7*). Moreover, Fe/S disruption and Fenton-mediated •OH production have been proposed as a downstream consequences of aminoglycoside killing (*30*). Given the elevated •OH level (FIG 4D; **Supplementary FIG S5C**) and increased susceptibility of a *dps*-deficient strain upon PA14 exposure to the AGXX®/Gm combination (FIG 5B), we wondered whether the increased ROS production impairs the activity of Fe/S cluster-containing enzymes. We prepared cell lysates from PA14 exposed to individual or combined AGXX® and Gm, and measured the activity of aconitase, an Fe/S-containing enzyme of the TCA cycle, whose activity depends on the presence of the Fe/S-cluster. While individual treatments with sublethal AGXX720C® or Gm concentrations had no impact on aconitase activity, exposure to a combination of AGXX720C® and Gm resulted in ∼75% loss in activity (FIG 6A). The loss in aconitase activity could be explained by the release of iron atoms, as it occurs during Fe/S cluster oxidation under O_2_^-^and H_2_O_2_ stress (*47, 48*). To test the role of free iron for the synergistic effect between AGXX720C® and aminoglycosides, we treated PA14 with the iron chelator 2’,2’ bipyridyl prior to their exposure to sublethal concentrations of AGXX720C® and Gm. Indeed, we found that PA14 survival was less impaired when cells had been treated with 2’,2’ bipyridyl prior to the presence of the AGXX720C®/Gm cocktail (FIG 6B). However, iron chelators, such as 2’,2’ bipyridyl, have been reported to reduce aminoglycoside killing in *E. coli,* a phenomenon we confirmed in *P. aeruginosa* **(Supplementary FIG S5D)**. Overall, our data suggest a potential role of free iron for the ROS-mediated toxicity of aminoglycoside exposure when in synergy with AGXX®.

**FIG 6.**
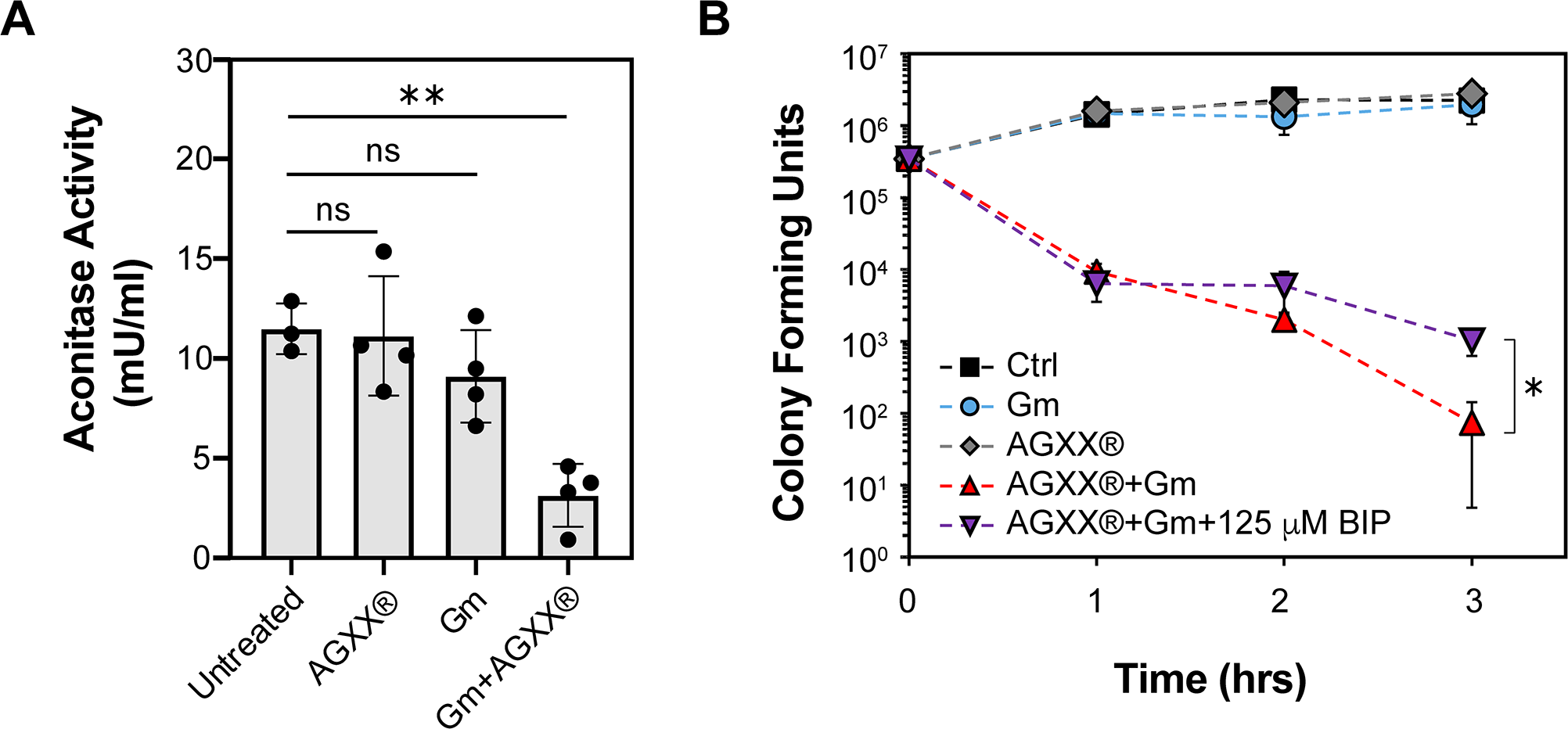
The synergy between AGXX® and aminoglycosides on PA14 killing is in parts mediated by a disruption in iron homeostasis. Overnight PA14 cultures were diluted into MOPSg and incubated under aerobic conditions until exponential phase was reached. **(A)** Cells were left untreated or treated with 100 µg/ml AGXX720C®, 0.6 µg/ml Gm, or the combination thereof for 1 hour. Aconitase activities were determined in crude extracts (*n=4, ± S.D*). One-way ANOVA, Dunnett’s posttest; *ns p>0.05,* ** *p* < 0.01. **(B)** Cultures were either left untreated (white square) or treated with 0.25 µg/ml Gm (white diamond), 50µg/ml AGXX720C® (white circle), or the combination thereof (white triangle) for three hours. Survival was determined each hour by serially diluting samples in PBS and plating onto LB agar for overnight growth. The impact of free iron on the increased killing by AGXX®/Gm cotreatments was tested by the absence (white triangle) and presence (black diamonds) of 125 µM 2’,2’ bipyridyl (*n=3, ± S.D*.). Student’s t test, * *p*<0.05).

### Combined AGXX® and aminoglycoside treatment induces significant membrane damage

In the initial stages of aminoglycoside uptake, the polycationic aminoglycoside electrostatically interacts with the bacterial outer membrane, displacing membrane divalent cations and inevitably increasing membrane permeability (*49*). Thus, compounds with membrane permeabilizing properties have been reported as potent aminoglycoside adjuvants (*50, 51*). To probe the role of membrane permeability, we evaluated outer membrane disruption using the hydrophobic fluorescent probe N-phenyl-1-napthylamine (NPN), which has diminished fluorescence in the presence of an intact outer membrane but significantly increases when bound to exposed phospholipid groups in a disrupted lipopolysaccharide monolayer (*52*). We found that in contrast to individual treatments with sublethal AGXX720C® and Gm concentrations, a combined treatment significantly increased NPN uptake, resulting in ∼7-fold higher NPN fluorescence (FIG 7A). Next, we examined the inner membrane permeability of PA14 cells that were subjected to AGXX720C® and Gm treatment alone and in combination, using the fluorescent probe propidium iodide (PI). Due to its size and charge, PI can only cross compromised inner membranes where it binds nonspecifically to nucleic acids enhancing its fluorescence exponentially (*53*). Individual treatments with sublethal AGXX720C® and gentamicin concentrations did not result in significantly increased PI fluorescence (FIG 7B) suggesting that at these concentrations none of them causes significant plasma membrane damage. A combined treatment, however, led to substantially increased PI fluorescence. Our spectrophotometric findings were complemented by fluorescent microscopy analyses of PA14 cells that were treated as described before, washed in PBS, stained with Syto9/PI (live/dead stain), incubated in the dark for 15 minutes at room temperature, mounted onto a glass slide with 1% agarose, and imaged at 63x magnification via inverted confocal microscopy. While the individual treatments with 0.25 µg/ml Gm or 50 µg/ml AGXX720C® only caused increased PI fluorescence in very few cells, the number of PI-stained cells was substantially higher in PA14 cells that were treated with a combination of sublethal AGXX720C®/Gm concentrations (FIG 7C).

**FIG 7.**
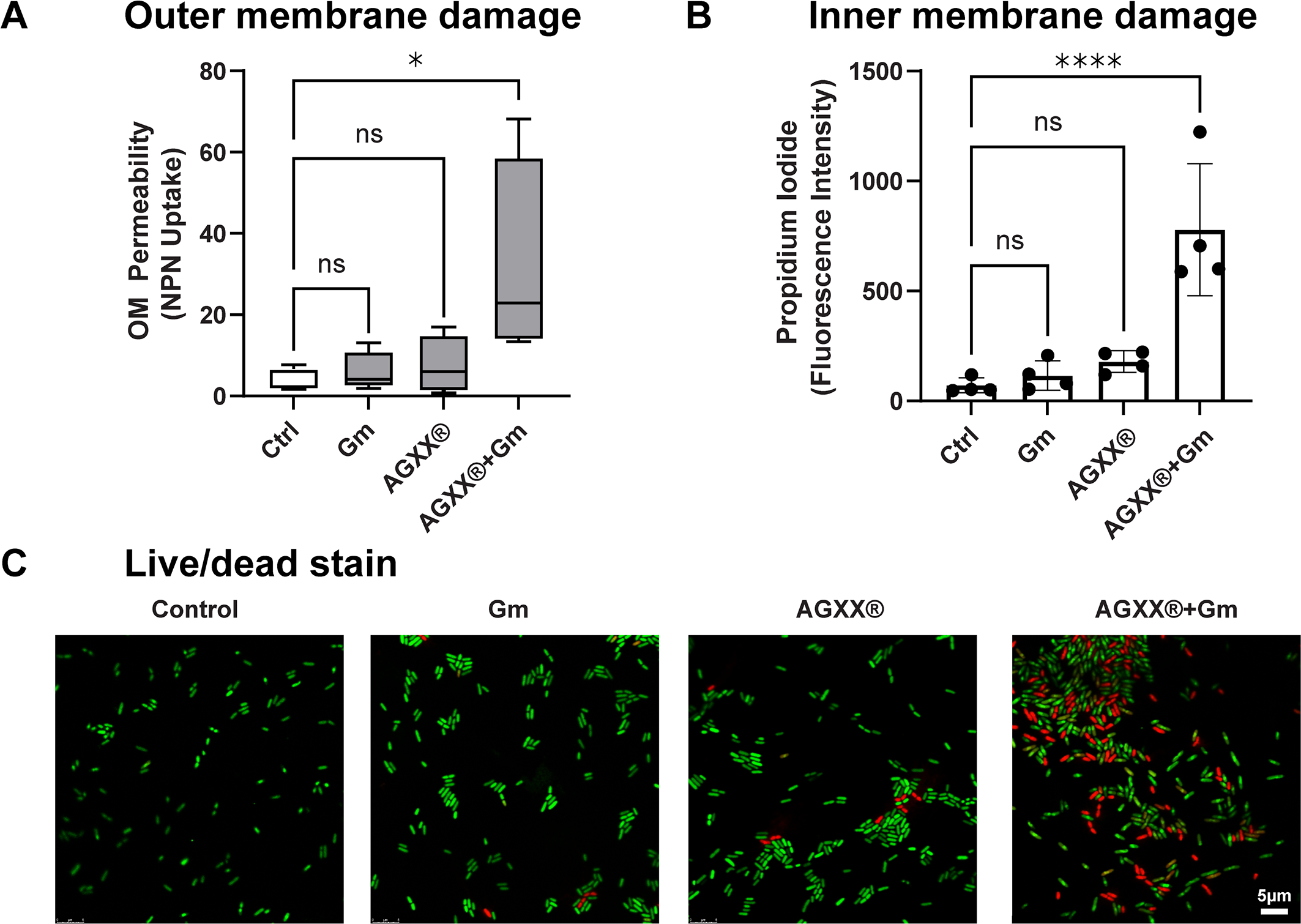
Combined AGXX® and aminoglycoside treatment induces significant membrane damage. PA14 cells grown to mid-log phase in MOPSg media were left untreated or treated with sublethal concentrations of Gm (0.25 µg/ml), AGXX720C® (50 µg/ml), and the combination thereof. Cells were harvested after 1 hour of treatment, washed in PBS, and stained with **(A)** 10 µM N-phenyl-1-naphthylamine (NPN) dye, and **(B)** 0.5 µM propidium iodide (PI). Fluorescence intensities were determined at excitation/emission wavelengths of **(A)** 350/420 nm and **(B)** 535/617 nm, respectively, (*n=4, ± S.D*.). One-way ANOVA, Dunnett’s posttest; *ns p>0.05,* * *p* < 0.05, **** *p* < 0.0001. **(C)** Samples were washed in PBS, stained with PI/Syto9, incubated in the dark for 15 minutes at room temperature, mounted onto a glass slide with 1% agarose, and imaged at 63x magnification via inverted confocal microscopy. One representative of three independent experiments with similar outcomes.

### AGXX® increases aminoglycoside uptake and lethality through increased activity of the proton motive force (PMF)

Considering the significance of membrane permeability for aminoglycoside uptake, we reasoned that the increased membrane disruption PA14 experiences during a combined treatment of AGXX® and aminoglycosides would facilitate intracellular accumulation of aminoglycosides. Using flow cytometry, we evaluated the uptake of Texas-Red-labelled Gentamicin (TR-Gm) in exponentially growing PA14 in the presence or absence of AGXX720C®. Treatment of PA14 with a combination of AGXX720C® and TR-Gm resulted in a significant increase in TR-Gm uptake similar to treatments with the membrane-targeting antibiotic polymyxin B (FIG 8A, **Supplementary FIG S6A).** The increased uptake of TR-Gm in the presence of AGXX® resulted in a 2.5 log reduction in bacterial survival compared to PA14 that were only exposed to TR-Gm alone **(Supplementary FIG S6B**). Moreover, the secondary and tertiary stage of aminoglycoside import appears to depend on an active membrane potential and occurs therefore only in actively respiring cells (*49, 54*). As such, increasing cellular respiration or stimulation of membrane potential has been found to increase aminoglycoside lethality (*55*). Using the protonophore CCCP, a compound that efficiently inhibits the proton motive force (PMF), we determined whether the synergy between AGXX® and aminoglycosides is dependent on the membrane potential. Not surprisingly, PA14 killing by a lethal dose of gentamicin (1 µg/ml) could be reduced by almost 2 logs when the cells were pre-incubated with CCCP (**Supplementary FIG S6C).** Likewise, pretreatment with CCCP restored PA14 survival to a similar degree in the presence of 0.25 µg/ml Gm and 50 µg/ml AGXX720C® for three hours (Fig. 8B). Surprisingly, combining polymyxin B at 1x MIC and 2x MIC with sublethal Gm concentrations did not result in increased killing relative to polymyxin B alone **(Fig. S2D)**. Overall, our findings indicate that AGXX®’s synergistic effect on aminoglycoside lethality does not only involve increased membrane permeability but also requires an active proton motive force across the bacterial membrane.

**FIG 8.**
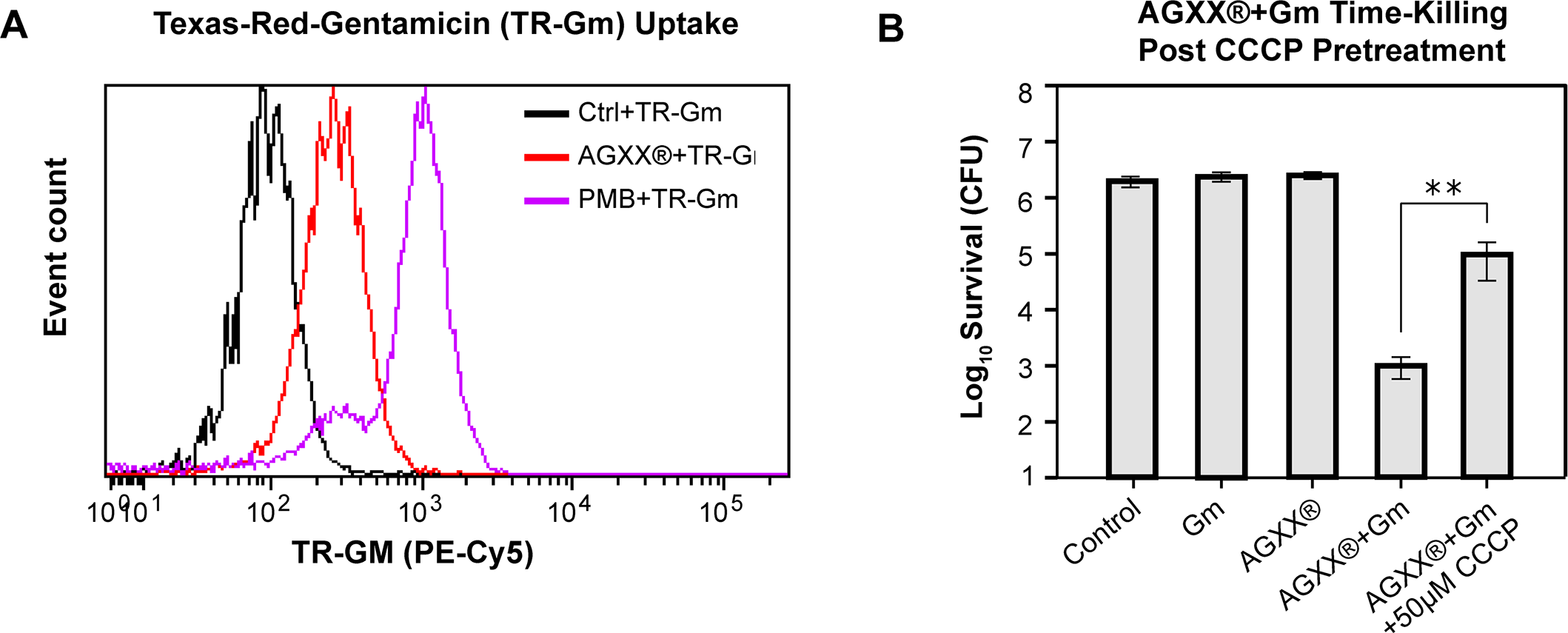
AGXX® increases aminoglycoside uptake and lethality through increased activity of the proton motive force (PMF) **(A)** Mid-log PA14 cells were treated with 1.0 µg/ml TR-Gm, 1.0 µg/ml TR-Gm + 50 µg/ml AGXX720C®, and 1.0 µg/ml TR-Gm + 2.0 µg/ml polymyxin B (PMB) for 1 hour, respectively. TR-Gm uptake was measured via flow cytometry. **(B)** Mid-log phase PA14 were left untreated (control) or exposed to 0.25 µg/ml Gm, 50 µg/ml AGXX720C® or 0.25 µg/ml Gm + 50 µg/ml AGXX720C® for 3 hours. Samples were serial diluted, plated on LB agar, and incubated for 20 hours for CFU counts. To test the impact of the PMF on the killing of a combination of AGXX® and aminoglycosides, PA14 were pretreated with or without 50 µM carbonyl cyanide *m*-chlorophenyl hydrazone (CCCP) prior to AGXX®/Gm exposure *(n=3, ± S.D.)*. Student’s t test, ** *p*<0.01.

## DISCUSSION

In the present study, we demonstrate evidence that the novel silver containing antimicrobial AGXX® may potentially be used an antibiotic adjuvant with activity on the opportunistic pathogen *P. aeruginosa*. We show that AGXX® serves as an efficient tool to potentiate the activity of aminoglycoside antibiotics by reducing bacterial survival up to 50,000-fold. Aside from a 1-log increase in killing found when the Ag/Ru complex was combined with the membrane-targeting antibiotic polymyxin B, AGXX® did not influence the activity of antibiotics targeting DNA replication, cell wall synthesis, or folate metabolism (FIG 2) raising the possibility that AGXX®’s potentiating effects could be linked to a malfunction in protein synthesis. AGXX®’s potentiating effect was not limited to select aminoglycosides but applied to all members tested, including gentamicin, streptomycin, amikacin, and tobramycin (FIG 3). We observed the exponential killing of the AGXX®/aminoglycoside combination in complex and minimal media independent of the presence or absence of amino acids, indicating that the media composition has only minor effects on the synergistic effect (**Supplementary FIG S3)**. Additional support for AGXX®’s stimulating effect on aminoglycoside activity was provided by its ability to re-sensitize a kanamycin resistant *P. aeruginosa* strain (FIG 4). Notably, both AGXX® and aminoglycoside antibiotics were present at concentrations far below their MICs (0.1-0.3x MIC) indicating that the combination between the compounds was quite powerful with regards to increasing the bactericidal activity of aminoglycosides.

Based on our findings, we propose the following model for the synergy between AGXX® and aminoglycoside antibiotics (FIG 9): The antimicrobial action of AGXX® is proposed to involve O_2_^-^ and H_2_O_2_, which are generated in a redox cycle between silver (Ag) and ruthenium (Ru^x+1^) (*15, 16*). While the location of AGXX®-mediated ROS production was not the focus of this study, we propose that ROS is mainly generated in the extracellular part before the compounds penetrate the cell. This assumption is based on our finding that exogenous addition of catalase was highly effective in reducing ROS level and increasing *P. aeruginosa* survival upon treatment with a combination of AGXX® and Gm (**Supplementary FIG S5A**), given that catalase cannot readily cross the bacterial cell envelope. The combination of AGXX® and aminoglycosides concertedly increased endogenous ROS levels, including H_2_O_2_ and •OH. Potentially facilitated by released silver ions, the increased ROS level may disrupt iron-sulfur clusters in metabolic enzymes such as aconitase, resulting in the release of free iron, which ultimately triggers •OH formation in a Fenton reaction (*56*). Increasing ROS levels can inflict macromolecule damage such as DNA damage, protein aggregation, and membrane damage, as evidenced by *(i)* the increased expression of members of the SOS and heat shock responses, and *(ii)* the possibility that the observed damage on outer and inner membrane is mediated by ROS. In either way, increased ROS production contributes to the enhanced killing of *P. aeruginosa* upon treatment with a combination of AGXX® and aminoglycosides. The synergistic effect between AGXX® and aminoglycosides is also mediated by an increased uptake and the cellular accumulation of aminoglycosides, which can be attenuated by disrupting the bacterial membrane potential with ionophores such as CCCP.

**FIG 9.**
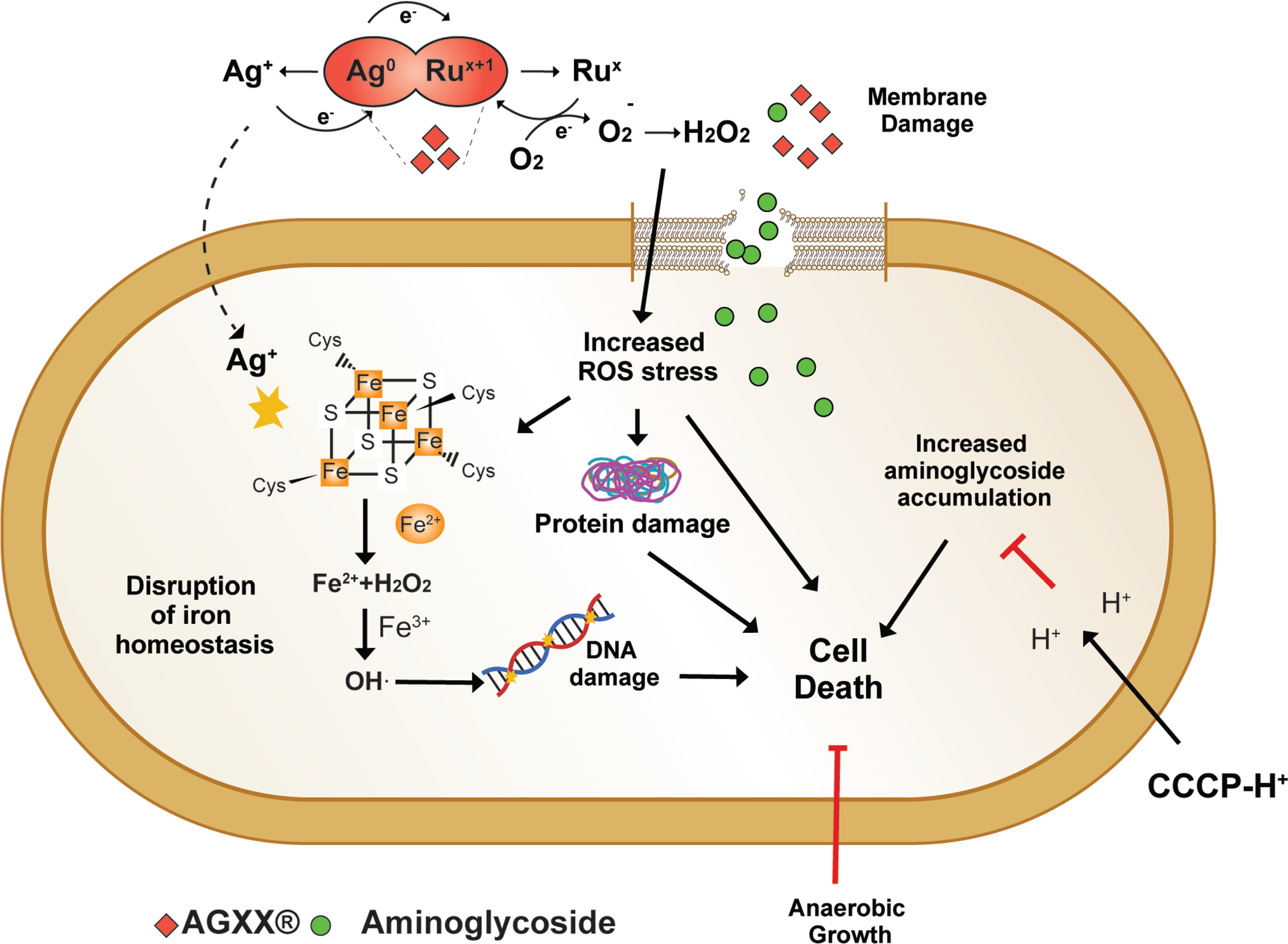
Proposed model for the synergy between AGXX® and aminoglycoside antibiotics. The antimicrobial action of AGXX® is mediated by superoxide (O_2_^-^) and hydrogen peroxide (H_2_O_2_), which are generated in a redox cycle between silver (Ag) and ruthenium (Ru^x+1^). The combination of AGXX® and aminoglycosides concertedly increases endogenous ROS levels. As such anaerobic growth suppressed the synergistic interaction between the two antimicrobials at lower Gm concentrations. Potentially facilitated by a release of silver ions, the increased ROS level may disrupt iron-sulfur clusters in metabolic enzymes such as aconitase, resulting in the release of free iron, which ultimately triggers hydroxyl radical (•OH) formation in a Fenton reaction. Increasing ROS levels can inflict macromolecule damage such as DNA damage and protein aggregation, contributing to increased killing as observed for *P. aeruginosa* that were treated with a combination of AGXX® and aminoglycosides. The synergistic effect between AGXX® and aminoglycosides is also mediated by an increased uptake and the cellular accumulation of aminoglycosides, which can be attenuated by disrupting the bacterial membrane potential with ionophores such as CCCP.

The antimicrobial effects of silver and silver containing agents have been extensively documented (*57–59*). However, in spite of the increasing use of silver in antimicrobial applications, its mechanism of action remains to be poorly understood (*24, 60, 61*). Silver along with other ROS inducing agents have been proposed to induce an array of multimodal cytotoxic events, including the mis-metalation of proteins, DNA damage, and imbalanced redox homeostasis, which ultimately disrupt multiple bacterial networks leading to cell death (*5, 17, 62*). Thus, we investigated whether AGXX®, through its multimodal effects as an ROS inducing compound, could increase that activity of conventional antibiotics against *P. aeruginosa*. AGXX® itself has been proposed to exert its antimicrobial action through the generation of ROS rather than the release of silver ions. AGXX®-mediated ROS production likely starts with the generation of O_2_^-^, which subsequently can be dismutated by cellular superoxide dismutases into hydrogen peroxide (*15, 63*). Likewise, aminoglycosides have also been posited to induce metabolic alterations that induce endogenous ROS levels (*64*). Growth of *P. aeruginosa* under anaerobic conditions provided a protective effect against the bactericidal effects of AGXX®/aminoglycoside killing (FIG 5A). Considering that the availability of nitrate as an alternative electron acceptor would facilitate PMF activity during anaerobic growth, a decrease in killing efficiency points towards a relevant role for ROS in the synergistic combination of the two compounds. At higher gentamicin concentrations, however, AGXX® sensitized *P. aeruginosa* to gentamicin (FIG 6A), which could be potentially explained with an increase in aminoglycoside uptake due to the high gentamicin availability.

We found a significant increase in hydrogen peroxide and •OH radical signals upon exposure of PA14 to a combination of AGXX® and aminoglycoside as evidenced by increased PO1 and HPF fluorescence (FIG 4) and decreased survival of transposon insertion strains defective in KatA and Dps (FIG 5B). Surprisingly, a transposon insertion into the *oxyR* gene, encoding the major hydrogen response regulator OxyR, did not show a significant difference in survival relative to the wildtype under combined AGXX®/aminoglycoside exposure (FIG 5B). However, this is consistent with a previous study on the aminoglycoside potentiating effects of silver nitrate, in which neither *oxyR-* deficient strains nor strains that constitutively express OxyR had significant effects on bacterial survival (*28*). On the other hand, in a different study overexpression of the H_2_O_2_ detoxifying gene, *ahpF,* attenuated the lethality and proteotoxic effects of aminoglycosides, providing evidence for the relevance of oxidative stress in aminoglycoside toxicity (*65*). Previous studies have demonstrated that pretreatment with antioxidant molecules such as thiourea reduced intracellular •OH radical levels and increased bacterial survival during aminoglycoside and silver stress (*17, 30, 66, 67*). Consistent with these findings, we made similar observations as the addition of either catalase or thiourea to AGXX®-aminoglycoside-treated PA14 resulted in a significant reduction in endogenous ROS level as well as bacterial killing relative to the combined treatment alone (FIG 4A, B; **Supplementary FIG S5A, B)**. Consistent with our previous report that AGXX® causes protein aggregation (*68*), we found significant transcriptional upregulation of *ibpA*, which encodes the molecular chaperone IbpA, during exposure to AGXX® and Gm. Given that IbpA serves as the front line defense to limit the aggregation of unfolded proteins (*44*), AGXX®/Gm-induced oxidative stress likely causes an imbalance of proteostasis and increases the cellular demand for chaperone expression. Interestingly, heat-shock mediated protein aggregation was found to exponentially increase aminoglycoside lethality in several gram-negative bacteria (*69*). Like our study, this novel mechanism was attenuated under ROS quenching conditions (*69*), providing additional evidence for the relevance of ROS in aminoglycoside killing. Considering that individual AGXX® or gentamicin treatments did not result in a significant increase in ROS signals, it is plausible that applying these antimicrobials at concentrations far below the MIC would not induce the physiological/metabolic alterations necessary for aggravating endogenous ROS levels.

Considering that both O_2_^-^ and silver ions have been reported to destabilize Fe/S cluster proteins resulting in the release of solvent-exposed iron (*10, 70*), we investigated the effects of individual and combinational treatments with AGXX® and/or aminoglycoside antibiotics on the activity of aconitase, an Fe-S cluster containing enzyme. We found aconitase to be extremely sensitive towards exposure to both antimicrobials when administered in combination (FIG 6A). Disrupted Fe-S clusters could potentially lead to an increase in intracellular iron levels and subsequent •OH generation through Fenton reaction as it has been demonstrated under hydrogen peroxide stress in ROS-sensitive *E. coli* strains (*71*). This would explain the significantly higher •OH levels detected in PA14 cells that were exposed to a combined AGXX® and gentamicin mixture (FIG 4D), which could cause significant DNA damage and induce the SOS response. In fact, we found that the combination of sublethal AGXX® and gentamicin significantly induced the expression of *sulA,* an SOS response marker of DNA damage (FIG 4E). Likewise, addition of 2’,2’ bipyridyl resulted in a 1-log increase in bacterial survival when cells were exposed to the toxic cocktail of gentamicin and AGXX® suggesting that free iron contributes to the bactericidal effects (FIG 6B). However, a similar protective effect was observed in the presence of lethal Gm concentrations, suggesting that the possibility that the rescuing effect in the presence of BIP could be due to its limiting effect on aminoglycoside uptake and not necessarily a quenching of hydroxyl radical generation through Fenton reactions. The Ezraty lab has demonstrated that bipyridyl inhibits Fe-S biosynthesis under iron limiting conditions, resulting in reduced respiratory complex I and II activities and decreased PMF, which is necessary for aminoglycoside uptake (*28*). All in all, our findings highlight the relevance of oxidative stress in the enhancement of antibiotic lethality in *P. aeruginosa* exposed to sub-inhibitory concentrations of AGXX® and aminoglycosides.

Compared to other gram-negative pathogens, *P. aeruginosa* possesses high intrinsic resistance mechanisms towards a variety of antibiotics (*72*). This resistance is mediated in part by its extensive arsenal of efflux pumps and significantly lower outer membrane permeability (*72*). We found that combining AGXX® and gentamicin increased both inner and outer membrane permeability in *P. aeruginosa* leading to a subsequent increase in aminoglycoside uptake (FIG 7, FIG 8). Aminoglycosides are polycationic in nature and known to displace divalent cations that cross-bridge lipopolysaccharides, which makes the outer membrane substantially more permeable (*73, 74*). It is therefore possible that AGXX®, much like aminoglycosides, disrupts the outer leaflet of the outer membrane, with the result that lower amounts of aminoglycosides are required to permeabilize *P. aeruginosa*. If this were true, we would suspect that the initial stages of aminoglycoside uptake could likely be accelerated causing a more rapid antibiotic influx. Surprisingly, we found that the addition of sublethal polymyxin B concentrations did not affect gentamicin lethality indicating that the mechanism behind the potentiating effects of AGXX® extends beyond increased membrane permeability **(Supplementary FIG S6D)**. We further found that disrupting membrane potential using an ionophore increased bacterial survival under combined AGXX® and gentamicin stress (FIG 8), suggesting an important role for the energy dependent phase of aminoglycoside uptake AGXX®’s potentiating effect, as has been found with strategies that increase aminoglycoside lethality via TCA cycle mediated PMF generation (*55*).

By definition, antibiotic adjuvants do not possess inherent antimicrobial activity (*75*). Even though AGXX® can technically not be defined as an antibiotic adjuvant given its own antimicrobial activity (*16*), our use of sublethal AGXX® concentrations to screen for potentiating effects satisfied this definition. Notably, the antimicrobial properties of silver have been accepted for a long time and taken advantage of in topical ointments such as silverdene cremes to prevent and treat *P. aeruginosa* infections in burn wound patients (*26*). A comparison to silver nitrate and silver sulfadiazine (silverdene) revealed an increased potency of some of the AGXX® formulations resulting in a lower survival rate of PA14 (FIG 1). Moreover, given that antibacterial combination therapies typically involve significantly lower concentrations of the antimicrobials compared to what is needed for individual treatments, such approaches would potentially delay resistance development.

## MATERIALS AND METHODS

### Bacterial strains and growth conditions

Unless stated otherwise, overnight *P. aeruginosa* PA14 strains **(Supp. Table 2)** were grown under aerobic conditions in Luria-Bertani broth (LB, Millipore Sigma) at 37°C for 16-20 hours at 300 rpm. For subsequent assays, overnight cultures were diluted into Mueller-Hinton broth (MHB) or 3-(N-morpholino) propanesulfonic acid minimal media containing 0.2% glucose, 1.32 mM K_2_HPO_4_ and 10 μM thiamine (MOPSg) and incubated at 37 °C at 150 rpm. Growth under anaerobic conditions was performed as described previously (*76*). In brief, PA14 was grown anaerobically in the presence of 1% KNO_3_ in a 50 ml falcon tube that was additionally sealed with parafilm. Growth was performed in DuoClick™ (Thomas Scientific) screw cap culture tubes filled with minimal room. Overnight PA14 cultures were diluted to an OD_600_=0.002 in MHB containing 1% KNO_3_ and grown until OD_600_ ∼0.2 after which the treatments were applied for four hours.

### Minimum Inhibitory Concentration (MIC)

MIC assays were performed in 96-well plates in a total volume of 200 μl per well. Overnight PA14 cultures were diluted into MHB to an OD_600_ of 0.002 and distributed into 96 well plates that contained increasing concentrations of ciprofloxacin, nalidixic acid, carbenicillin, imipenem, trimethoprim, polymyxin B, gentamicin, amikacin, tobramycin, streptomycin, and kanamycin, respectively. Plates were subsequently incubated at 37 °C for 16 hours at 300 rpm. MIC assays were performed in duplicates. The MIC is defined as the lowest antibiotic concentration that inhibited growth.

### Time Killing Assays

Overnight PA14 cultures were diluted ∼25-fold into MHB by normalizing cultures to an OD_600_ = 0.1 in a 6-well sterile cell culture plate. Sublethal concentrations of AGXX720C® and antibiotics of interest were added individually and in combination as indicated and plates incubated at 37°C for 3 hours at 150 rpm under aerobic conditions. For thiourea, CCCP and BIP pretreatments, overnight PA14 was grown in MOPSg to exponential phase. Thiourea, CCCP or BIP was then added 60 minutes prior to AGXX®/aminoglycoside treatments whenever indicated. At 1-hour intervals, OD_600_ of cultures were recorded, sample volumes were normalized to the lowest optical density measured, serially diluted in PBS (pH = 7.4) and plated on LB agar to quantify colony forming units (CFU) after 16 hours incubation. Percent survival was calculated as the ratio of surviving colonies from treated to untreated samples.

### Intracellular ROS level

Intracellular ROS levels were quantified using the redox-sensitive dye 2’,7’-dichlorodihydrofluorescein diacetate (H_2_DCFDA) (Thermofisher Scientific). Exponentially growing PA14 cultures were left untreated or treated with 0.25 µg/ml Gm, 50 µg/ml AGXX720C® or a combination of both for 1 hour. Samples were collected and normalized to an OD_600_=1.0. Cells were washed twice, resuspended in prewarmed PBS containing 10 µM H_2_DCFDA, and incubated in the dark at 37 °C for 30 min before samples were washed twice again in PBS and DCF fluorescence measured at excitation/emission wavelengths of 485/535 nm (Tecan 200 plate reader). To quench cellular ROS, cells were pretreated with 50 mM thiourea prior to the stress treatments.

### Quantification of hydrogen peroxide level

The monoborate fluorescent probe peroxy orange 1 (PO1) was used to measure hydrogen peroxide level. Exponentially growing PA14 cultures were treated with 10 µM PO1 prior to their exposure to 0.25 µg/ml Gm, 50 µg/ml AGXX720C®, or a combination of both for 1 hour. Cells were washed twice and resuspended in prewarmed PBS. Samples were analyzed using the flow cytometer (BD FACS Melody™) in the PE-CF594 channel. At least 10,000 events were recorded, and figures generated using FCSalyzer (*77*).

### Quantification of Hydroxyl radical level

The fluorescent probe hydroxyphenyl fluorescein (HPF; Invitrogen) was used to determine the amount of cellular hydroxyl radicals (•OH) produced (*33*). Exponentially growing PA14 cultures were either left untreated or treated for 1 hour with 0.25 µg/ml Gm, 50 µg/ml AGXX720C®, or the combination thereof. Samples were collected and normalized to an OD_600nm_ of 1.0 after washing twice with PBS (pH 7.4). Cells were then stained with 10 µM HPF and incubated in the dark for 30 minutes at 37 °C. Cells were washed twice and resuspended in prewarmed PBS. Samples were analyzed using the flow cytometer (BD FACS Melody™) in the FITC channel. At least 10,000 events were recorded, and figures generated using FCSalyzer (*77*).

### Gene expression analyses by qRT-PCR

Overnight PA14 cultures were diluted into MOPSg media to an OD_600_=0.1 and grown to mid-log phase (OD_600nm_=0.3). Cultures were left untreated or treated with 0.25 µg/ml Gm, 50 µg/ml AGXX720C®, or the combination thereof for 60 minutes. Transcription was stopped by the addition of an equal volume of ice-cold methanol. Total RNA was extracted from three biological replicates using a commercially available RNA extraction kit (Macherey & Nagel). Remaining DNA was removed using the TURBO DNA-free kit (Thermo Scientific). mRNA was reverse transcribed into cDNA using the PrimeScript cDNA synthesis kit (Takara). The following primer pairs were used for gene amplification: *ibpA* - TTCCGTCATTCCGTAGG/ AGGTCTTCTTCCTGG ; *sulA -* ACTGTTCCAGGAAGCGTTCT/ AGCGAAAGTTCGCTAAAGGC; *rrsD -* TATCAGATGAGCCTAGGTCGGATTA/ TTTACAATCCGAAGACCTTCTTCAC. qRT-PCRs were set up according to the manufacturer’s instructions (Alkali Scientific). Transcript levels of the indicated genes were normalized against transcript levels of the 16S rRNA-encoding *rrsD* gene and relative fold changes in gene expression were calculated using the 2^-ΔΔCT^ method(*78*).

### Aconitase Assay

Exponentially growing PA14 were treated with sublethal concentrations of AGXX720C® (100 μg/ml), Gm (0.6 μg/ml) or the combination thereof. Aconitase activity was measured from cell lysates using the Aconitase Assay Kit (Abcam) according to manufacturer’s instructions. One unit of aconitase is defined as the amount of enzyme that isomerizes 1 μmol of citrate to isocitrate per min at pH 7.4 and 25 °C.

### Outer membrane permeability assay

The N-phenyl-1-naphthylamine (NPN) uptake assay was used to detect outer membrane damage as described in (*79*). Exponential phase PA14 were treated with 0.25 µg/ml Gm, 50 µg/ml AGXX720C®, a combination thereof. Addition of 2 μg/ml polymyxin B was used as a positive control. Cells were collected after 60 min of treatment, washed, and resuspended to an OD_600_=0.5 in HEPES-sodium buffer (pH 7.2). 10 µM NPN was added, and samples incubated in the dark for 15 min. NPN fluorescence was measured at excitation/emission wavelengths of 350/420 nm. Increased NPN uptake (indicating outer membrane damage) was calculated using the following equation (*80*):

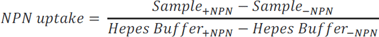

### Inner membrane disruption assay

Inner membrane integrity was determined by the cellular uptake of propidium iodide following antimicrobial treatments. Exponentially growing PA14 cells were either left untreated or treated with 0.25 µg/ml Gm, 50 µg/ml AGXX720C®, or a combination thereof. Cells were harvested after 1 hour, washed twice, and resuspended in PBS (pH 7.4) at an OD_600_=0.5. Propidium iodide (Thermo Fisher Scientific) was added to a final concentration of 500 nM and samples incubated in the dark for 30 min. Fluorescence intensities were measured at excitation/emission wavelengths of 535/617 nm. Samples treated with polymyxin B at a sublethal concentration (2 μg/ml) were included as a positive control.

### LIVE/DEAD staining

Exponentially growing PA14 cells were either left untreated or treated with 0.25 µg/ml Gm, 50 µg/ml AGXX720C®, or a combination thereof. Cells were harvested after 1 hour, washed twice, and resuspended in PBS (pH 7.4) at an OD_600_=0.2. Samples were stained with SYTO9 (6µM) and PI (30 µM) and incubated in the dark for 15 min at room temperature. Cells were then transferred onto a glass slide and covered with 1% agarose prior to visualization using a Leica SP8 confocal system equipped with a DMi8 CS inverted microscope. Samples treated with polymyxin B at a sublethal concentration (2 μg/ml) were included as a positive control.

### Texas Red-Gentamicin Uptake Assay

Texas Red-succinimidyl ester (Invitrogen) was dissolved to a final concentration of 20 mg/ml in high quality anhydrous N,N-dimethylformamide at 4 °C (*81*). Gm was dissolved in 100 mM K_2_CO_3_ (pH 8.5) to a final concentration of 10 mg/ml at 4 °C. 20 µl Texas Red was slowly added dropwise to 700 ul Gm to allow for the conjugation reaction to obtain the Texas-Red labeled gentamicin (TR-Gm) (*81*). The TR-Gm conjugate was diluted in water and stored at −20 °C protected from light. Exponentially growing PA14 cells were first treated with 1 µg/ml TR-Gm followed by exposure to either 50 µg/ml AGXX720C® or 2 µg/ml polymyxin B (positive control). After 1 hour incubation in the dark at 37 °C, cells were collected and analyzed in the flow cytometer (BD FACS Melody™) using the PE-Cy5 channel. At least 10,000 events were recorded, and figures generated using FCSalyzer (*77*).

### Statistical analyses

All statistical analyses were performed in GraphPad prism version 8.0.

## ACKNOWLEDGEMENTS

This work was supported by the NIAID grant R15AI164585 and the Illinois State University Pre-Tenure Faculty Initiative Grant (to J.-U. D.). G.Y.D. was supported by Weigel and Mockford-Thompson by the Phi-Sigma Biological Sciences Honors Society, and the SIGMA Xi and BIRDFeeder grants. G.M.A. and C.D.O. were supported by the Illinois State University Undergraduate Research Support Program. Dr. Uwe Landau and his team from Largentec GmbH are acknowledged for scientific discussion and providing the AGXX® formulations. We further thank the ISU Confocal Microscopy Facility which was funded by NSF grant DBI-1828136, for their support. We also thank Sadia Sultana for critically proof-reading the manuscript and providing invaluable feedback. Figure 9 was generated with Adobe Illustrator.

