## Supplemental Fig S1-6 for "A Novel Silver-Containing Antimicrobial potentiates aminoglycoside activity against *Pseudomonas aeruginosa*"

### Supplementary Information

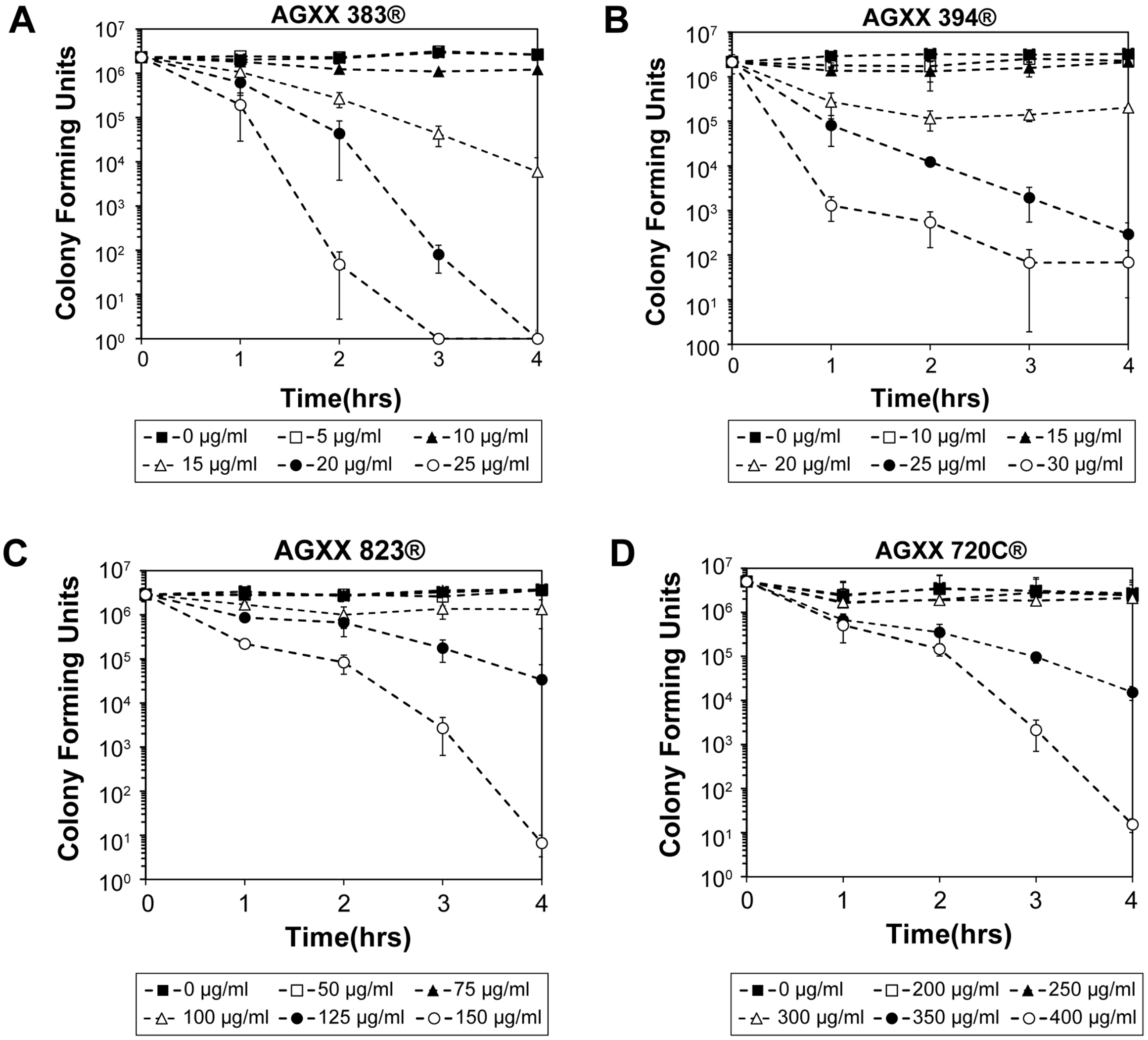


**FIG S1** PA14 cultures grown overnight in LB media were diluted 25-fold into MOPSg media and either left untreated (black square) or exposed to the indicated concentrations of **(A)** AGXX383®, **(B)** AGXX394®, **(C)** AGXX823®, and **(D)** AGXX720C®, respectively. For the course of four hours, samples were taken every 60 min, serial diluted, plated on LB agar, and incubated for 20 hours for CFU counts (*n=3, ± S.D.).*

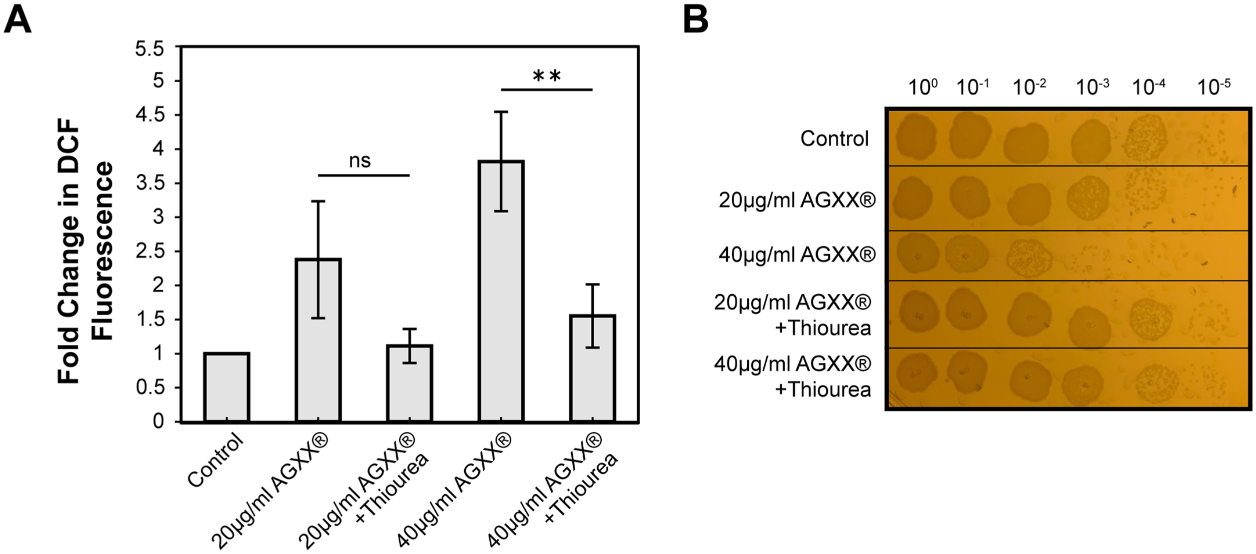


**FIG S2 AGXX® treatment elicits increased ROS levels in *P. aeruginosa*.** PA14 cells grown to mid-log phase were treated with either 20 µg/ml (sublethal) or 40 µg/ml (lethal) AGXX394® in MOPSg for 1 hour. **(A)** Intracellular ROS levels were quantified by $H_2$DCFDA fluorescence. 50 mM thiourea was used as a ROS quencher (*n=3, ± S.D)*. **(B)** PA14 cells were harvested after 1 hour of exposure, serially diluted in PBS and plated on LB agar to enumerate surviving colonies. Shown is one representative of three biological replicates.

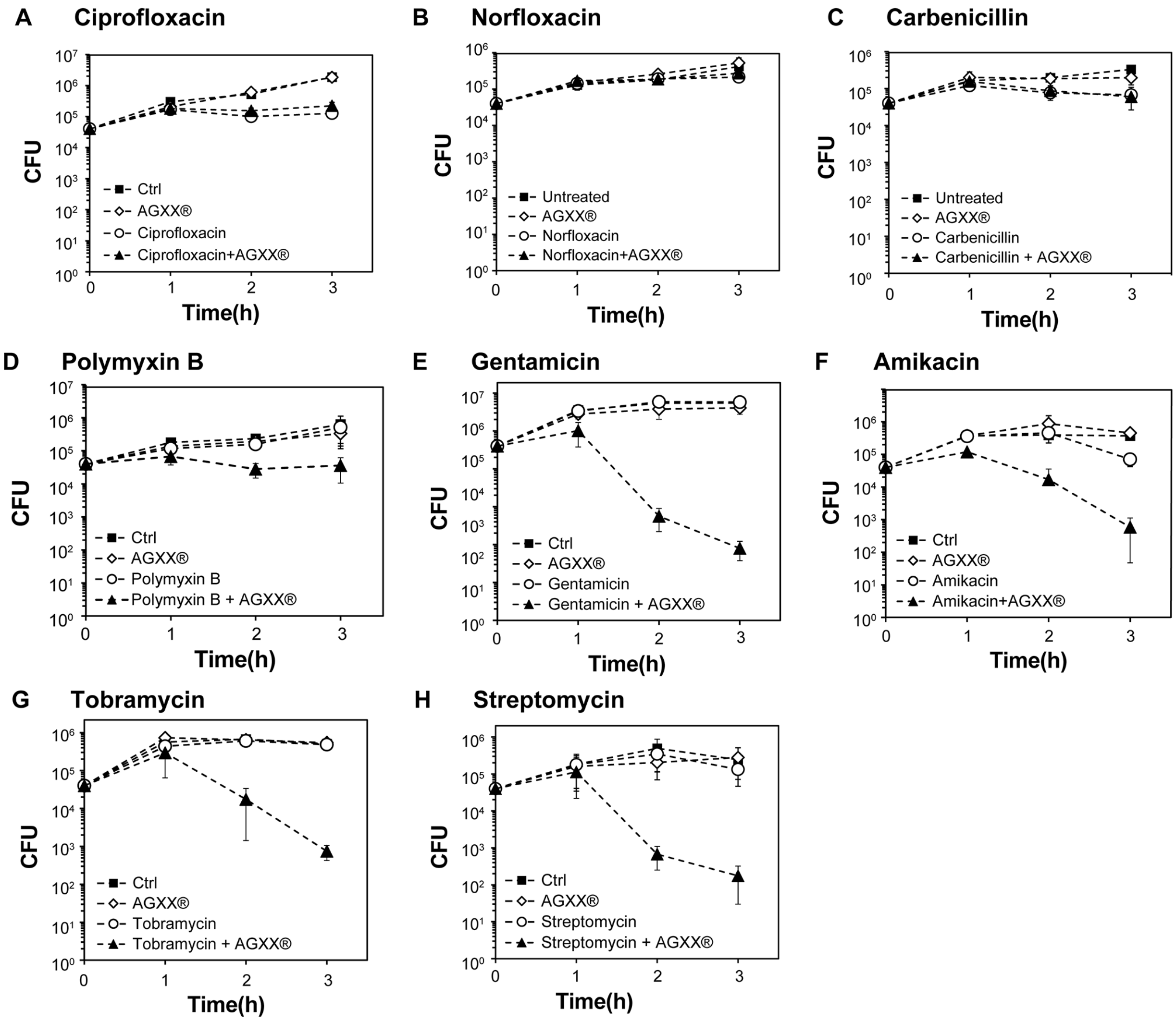


**FIG S3 AGXX® potentiates aminoglycoside activity against *P. aeruginosa.*** PA14 cultures grown overnight in LB media were diluted ~25-fold ($OD_{600}$=0.1) into MHB and either left untreated (black squares) or exposed to 75 µg/ml AGXX720C® (white diamonds), a sublethal concentration of the indicated antibiotic (white circles), or the combination thereof (black triangles). Over the course of 3 hours, samples were taken every 60 min, serial diluted, plated on LB agar, and incubated for 20 hours for CFU counts (*n=3, ± S.D.).*

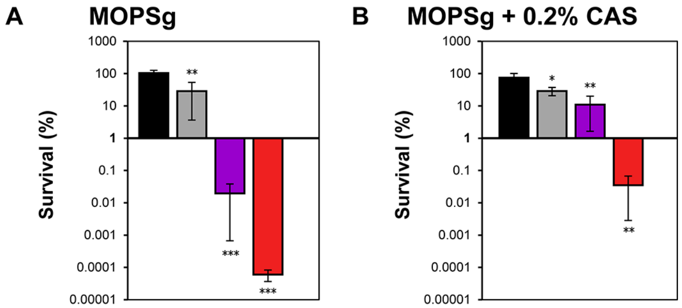


**FIG S4 The synergy between AGXX® and aminoglycosides is not media-specific.** Overnight PA14 cultures were diluted to an $OD_{600}$=0.05 in **(A)** MOPSg alone or **(B)** MOPSg supplemented with 0.2% casamino acids. Cultures were then grown until early log phase ($OD_{600}$=0.2) and treated with 0.3 µg/ml Gm (black bars), 0.3 µg/ml Gm + 10 µg/ml AGXX® (grey bars), 0.3 µg/ml Gm + 20 µg/ml AGXX® (violet bars), 0.3 µg/ml Gm + 30 µg/ml AGXX® (red bars) for four hours. PA14 cells were then harvested, serially, diluted in PBS and plated on LB agar to enumerate surviving colonies. (*n=3,* ± *S.D.*; One-way ANOVA, Dunnett's posttest relative to gentamicin treated sample; *ns*=$p>0.05$, * $p < 0.05$, ** $p < 0.01$, *** $p < 0.001$).

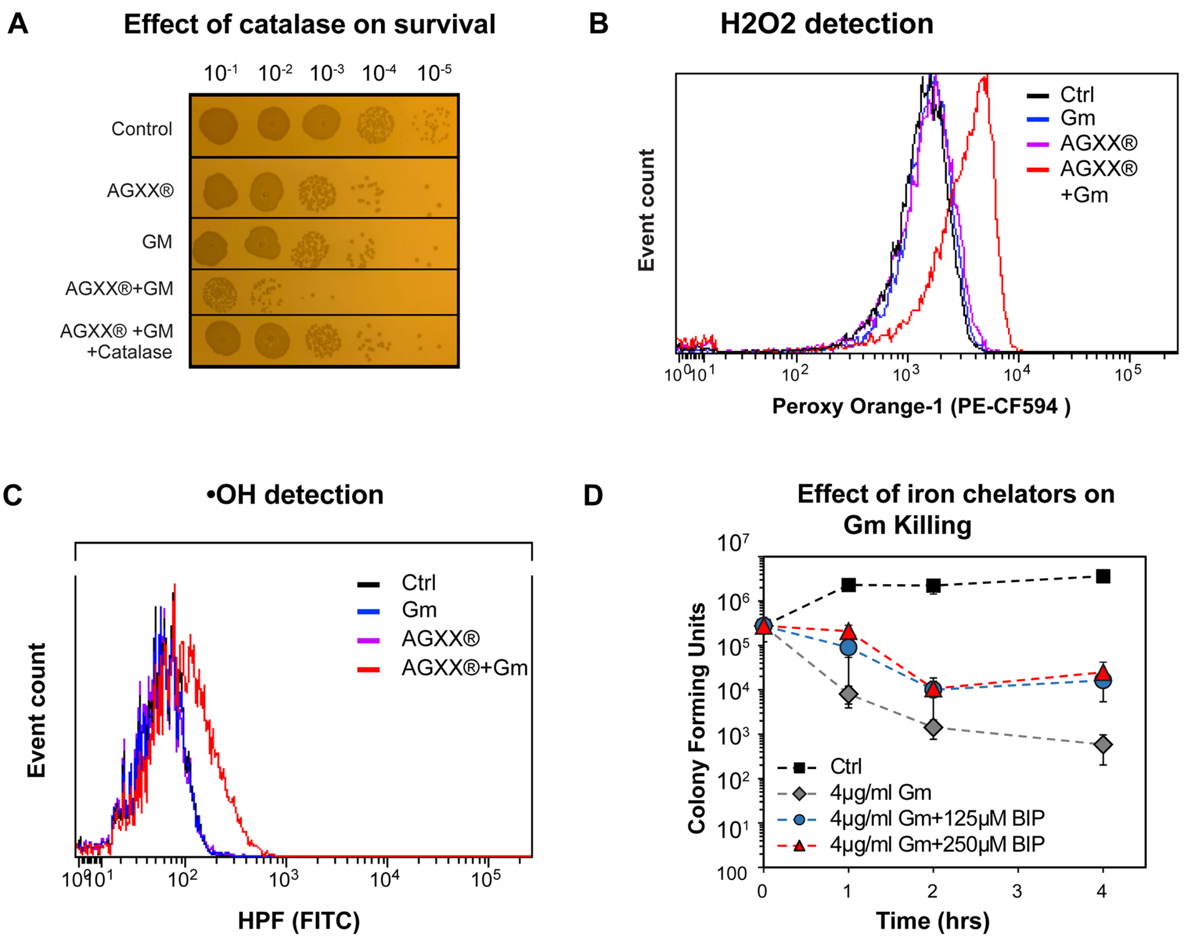


**FIG S5 Combined treatment with AGXX® and gentamicin increases endogenous ROS levels. (A)** Mid-log PA14 cells were treated with sublethal concentrations of Gm (0.25 µg/ml), AGXX720C® (50 µg/ml), the combination thereof, or left untreated. Preexposure to catalase was used as a ROS quencher. Samples were serially diluted in PBS after 60 min of incubation, spot-titered onto LB agar and incubated for 20 hours. One representative of three independent experiments with similar outcomes. **(B, C)** Exponentially growing PA14 cells were either left untreated or treated with sublethal concentrations of Gm (0.5 µg/ml), AGXX720C® (50 µg/ml), and the combination thereof for 60 minutes. Intracellular ROS levels were quantified by staining PA14 cells with **(B)** 10 µM PO1 and **(C)** 10 µM HPF, respectively. Dye fluorescence was measured via flow cytometry after 30 min of incubation. One representative of three independent experiments of similar outcomes. **(D)** Mid-exponential phase PA14 ($OD_{600}$=0.3)

untreated/pretreated with 125μM or 250 μM 2’, 2’ bipyridyl was exposed to 4 μg/ml Gm. Survival was quantified for four hours by serially diluting in PBS and plating on LB agar (*n=3, ± S.D.)*.

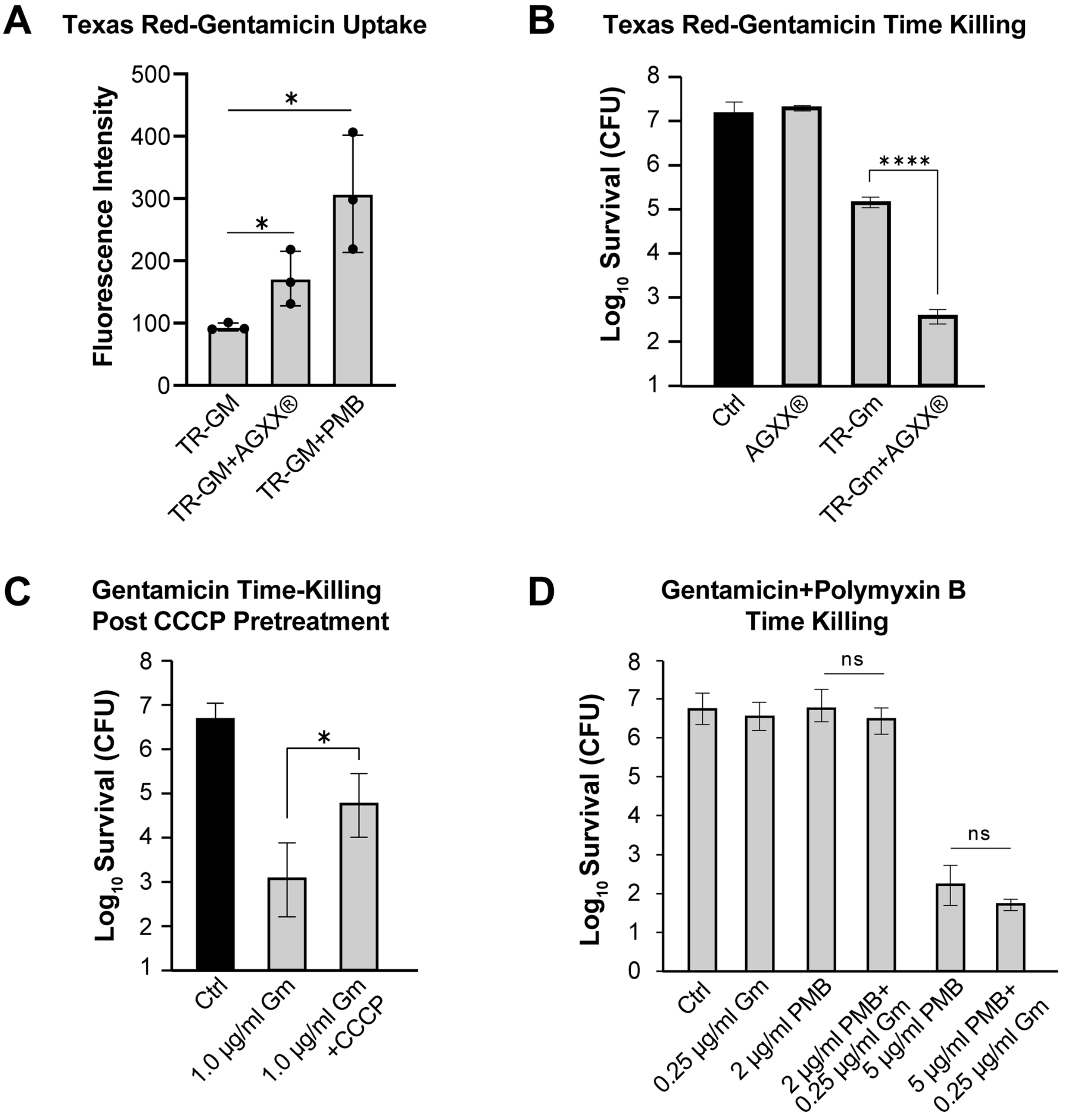


**FIG S6 AGXX® increases aminoglycoside uptake and lethality through increased PMF activity.** PA14 cells were grown to mid-log phase ($OD_{600}$=0.3) and treated with: **(A)** 1.0 µg/ml TR-Gm in the presence and absence of 50 µg/ml AGXX720C® and 2.0 µg/ml polymyxin B, respectively, to measure TR-Gm uptake by flow cytometry after 1 hour of treatment (*n=3, ± S.D., )*; (**B**) 50 µg/ml AGXX720C®, and 1.0 µg/ml TR-Gm in the

presence and absence of 50 µg/ml AGXX720C®. Samples were taken after 2 hours, serial diluted and plated on LB agar. CFU were counted after 20 hours ($n=3$, *± S.D.);* **(C)** To test the impact of the PMF on the killing of 1 µg/ml Gm, cells were pre-treated with 10 µM carbonyl cyanide *m*-chlorophenyl hydrazone (CCCP) prior to Gm exposure ($n=3$, *± S.D.)*; (**D**) polymyxin B (2 µg/ml and 5 µg/ml, respectively) with or without 0.25 µg/ml Gm for 3 hours. Samples were serial diluted, plated on LB agar, and incubated for 20 hours for CFU counts ($n=3$, *± S.D.)*. (Student's t test, ns $p>0.05$, * $p<0.05$, **** $p<0.0001$).
$p<0.05$, **** $p<0.0001$).
